# Evolution and global transmission of a multidrug-resistant, community-associated MRSA lineage from the Indian subcontinent

**DOI:** 10.1101/233395

**Authors:** Eike J. Steinig, Sebastian Duchene, D. Ashley Robinson, Stefan Monecke, Maho Yokoyama, Maisem Laabei, Peter Slickers, Patiyan Andersson, Deborah Williamson, Angela Kearns, Richard Goering, Elizabeth Dickson, Ralf Ehricht, Margaret Ip, Mathew V.N. O’Sullivan, Geoffrey W. Coombs, Andreas Petersen, Grainne Brennan, Anna C Shore, David C. Coleman, Annalisa Pantosti, Herminia de Lencastre, Henrik Westh, Nobumichi Kobayashi, Helen Heffernan, Birgit Strommenger, Franziska Layer, Stefan Weber, Hege Aamot, Leila Skakni, Sharon J. Peacock, Derek Sarovich, Simon Harris, Julian Parkhill, Ruth C. Massey, Mathew T.G. Holden, Stephen D. Bentley, Steven Y.C. Tong

## Abstract

The evolution and global transmission of antimicrobial resistance has been well documented in Gram-negative bacteria and healthcare-associated epidemic pathogens, often emerging from regions with heavy antimicrobial use. However, the degree to which similar processes occur with Gram-positive bacteria in the community setting is less well understood. Here, we trace the recent origins and global spread of a multidrug resistant, community-associated *Staphylococcus aureus* lineage from the Indian subcontinent, the Bengal Bay clone (ST772). We generated whole genome sequence data of 340 isolates from 14 countries, including the first isolates from Bangladesh and India, to reconstruct the evolutionary history and genomic epidemiology of the lineage. Our data shows that the clone emerged on the Indian subcontinent in the early 1970s and disseminated rapidly in the 1990s. Short-term outbreaks in community and healthcare settings occurred following intercontinental transmission, typically associated with travel and family contacts on the subcontinent, but ongoing endemic transmission was uncommon. Acquisition of a multidrug resistance integrated plasmid was instrumental in the divergence of a single dominant and globally disseminated clade in the early 1990s. Phenotypic data on biofilm, growth and toxicity point to antimicrobial resistance as the driving force in the evolution of ST772. The Bengal Bay clone therefore combines the multidrug resistance of traditional healthcare-associated clones with the epidemiological transmission of community-associated MRSA. Our study demonstrates the importance of whole genome sequencing for tracking the evolution of emerging and resistant pathogens. It provides a critical framework for ongoing surveillance of the clone on the Indian subcontinent and elsewhere.

**Importance:** The Bengal Bay clone (ST772) is a community-acquired and multidrug-resistant *Staphylococcus aureus* lineage first isolated from Bangladesh and India in 2004. In this study, we show that the Bengal Bay clone emerged from a virulent progenitor circulating on the Indian subcontinent. Its subsequent global transmission was associated with travel or family contact in the region. ST772 progressively acquired specific resistance elements at limited cost to its fitness and continues to be exported globally resulting in small-scale community and healthcare outbreaks. The Bengal Bay clone therefore combines the virulence potential and epidemiology of community-associated clones with the multidrug-resistance of healthcare-associated *S. aureus* lineages. This study demonstrates the importance of whole genome sequencing for the surveillance of highly antibiotic resistant pathogens, which may emerge in the community setting of regions with poor antibiotic stewardship and rapidly spread into hospitals and communities across the world.

## Introduction

Methicillin-resistant *Staphylococcus aureus* (MRSA) is a major human pathogen with a propensity to develop antibiotic resistance, complicating treatment and allowing persistence in environments where there is antibiotic selection pressure. While multidrug resistance has traditionally been the domain of healthcare-associated strains, the emergence of strains in the community setting that are also resistant to multiple antibiotics poses a significant challenge to infection control and public health (Tong and Kearns 2013). Given the heavy burden and costs associated with MRSA infections (Suaya et al. 2014; Tong et al. 2015), there is an urgent need to elucidate the patterns and drivers behind the emergence of drug-resistant community-associated MRSA lineages.

Over the last few years, several population genomic studies have started to unravel the evolutionary history of community-associated *S. aureus* lineages emerging in specific regions of the world. The prototype of these clones is the diverse USA300 lineage (ST8), forming distinct genetic lineages in North America and South America (Planet et al. 2013; Planet et al. 2015), including distinct clades in Europe and Africa (Strauß et al. 2017). The East-Asia clone (ST59) has diverged into two distinct lineages, with evidence of establishment in Taiwan and North America (Ward et al. 2016), while the ST80 lineage originated in North Africa, but went through a notable population expansion to become the dominant community-associated lineage in North Africa, the Middle East and Europe (Stegger et al. 2014). On the Australian continent, the Queensland clone (ST93) emerged in Indigenous communities of Western Australia and the Northern Territory, spread to the eastern seaboard and sporadically overseas, but forms a clade associated with Pacific Islander and Maori populations in New Zealand (van Hal et al. 2018).

This diversity of regional evolutionary histories is reflected in the various factors that have been suggested to contribute to the emergence and establishment of community-associated clones. For instance, acquisition of Panton-Valentine leucocidin (PVL), a mutation in the capsule gene *cap5D* and acquisition of the SCC*mec*-IV and ACME / COMER elements played a defining role in the regional evolution of USA300 (Strauß et al. 2017). While acquisition of a typical SCC*mec*-IV element was also associated with the emergence of the eastern seaboard clade of the Queensland clone, it appears that host population factors similar to those associated with the emergence in Indigenous populations of Australia were a possible driver in the establishment of Pacific Islander associated clade in New Zealand (van Hal et al. 2018). In contrast, the population expansion of the ST80 lineage was linked to the acquisition of a SCC*mec*-IV and fusidic acid resistance, and enrichment of resistance determinants in the Taiwan clade suggest a strong contribution for its emergence and persistence in East-Asia. While there is some evidence from these studies that resistance acquisition can be a driving force behind the regional emergence of community-associated MRSA clones, there is a lack of data on strains from critical regions that are considered hot-spots for the emergence of multidrug resistant pathogens, such as the Indian subcontinent.

In 2004, a novel *S. aureus* clone designated sequence type (ST) 772 was isolated from hospitals in Bangladesh (Afroz et al. 2008) and from a community-setting study in India (Goering et al. 2008). The clone continued to be reported in community- and healthcare-associated environments in India, where it has become one of the dominant epidemic lineages of community-associated MRSA (Chen and Huang 2014). Similar to other *S. aureus*, ST772 primarily causes skin and soft tissue infections, but more severe manifestations such as bacteraemia and necrotising pneumonia have been observed. Its potential for infiltration into nosocomial environments (D’Souza et al. 2010; Manoharan et al. 2012; Nadig et al. 2012; Blomfeldt et al. 2017) and resistance to multiple classes of commonly used antibiotics (including aminoglycosides, β-lactams, fluoroquinolones, macrolides and trimethoprim) (Chakrakodi et al. 2014; Steinig et al. 2015; Blomfeldt et al. 2017) has made ST772 an alarming public health concern on the Indian subcontinent and elsewhere. Over the last decade, the clone has been isolated from community- and hospital-environments in Asia, Australasia, Africa, the Middle East and Europe (SI Map 1, SI Table 1). As a consequence of its discovery, distribution and epidemiology, the lineage has been informally dubbed the “Bengal Bay clone” (Ellington et al. 2010). Despite clinical and epidemiological hints for a recent and widespread dissemination of ST772, a unified perspective on the global evolutionary history and emergence of the clone is lacking.

Here, we analyse whole genome sequences from a globally representative collection of 340 ST772 strains to elucidate the key events associated with the emergence and global spread of a multidrug resistant community-associated MRSA clone. Our analysis suggests that the clone originated on the Indian subcontinent in the 1970s and rapidly expanded through the 1990s and early 2000s. We found that international travel and family connections (India, Bangladesh, Nepal and Pakistan) in the region were closely linked with the global spread of the lineage. Genome integration of a multidrug resistance plasmid appeared to be a driver in the emergence of a dominant clade (ST772-A) in the early 1990s.

## Results

We generated whole genome sequence data of 354 *S. aureus* ST772 isolates collected across Australasia, South Asia, Hong Kong, the Middle East and Europe between 2004 and 2013 (SI Map 2, SI Table 2). Fourteen isolates were excluded after initial quality control due to contamination (SI Tables 2, 3). The remainder mapped with 165x average coverage against the PacBio reference genome DAR4145 (Steinig et al. 2015) from Mumbai (SI Tables 2, 3). Phylogenetic analysis using core-genome SNPs (n = 7,063) revealed little geographic structure within the lineage (Fig. 1a). Eleven ST772 methicillin-susceptible *S. aureus* (MSSA) and MRSA strains were basal to a single globally distributed clade (ST772-A, n = 329) that harbored an integrated resistance plasmid (IRP) described in the reference genome DAR4145 (Steinig et al. 2015) (Figs. 1a, 1b). Population network analysis distinguished three distinct subgroups within ST772-A (Figs. 1a, 1c): an early-branching subgroup harboring multiple subtypes of the staphylococcal chromosome cassette (SCC*mec*-V) (A1, n = 81), a dominant subgroup (A2, n = 153) and an emerging subgroup (A3, n = 56), that both exclusively harbor a short variant of SCC*mec*-V.

**Fig. 1:**
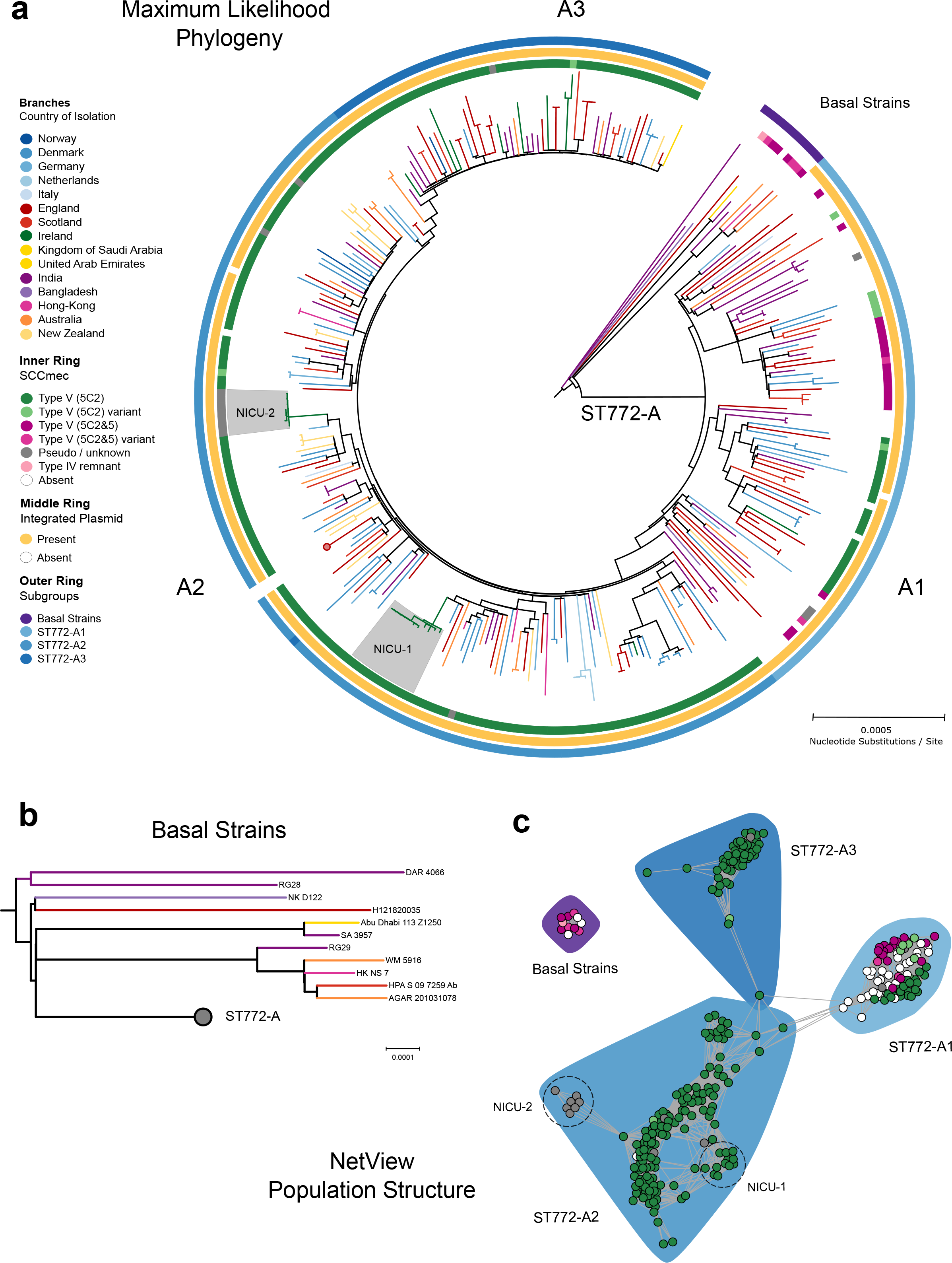
Evolutionary history and population structure of ST772. **(a)** Maximum likelihood phylogeny of ST772 (n = 340) based on 7,063 core-genome SNPs. Branch colors denote country of isolation, the inner ring delineates presence and type of SCC*mec*, the middle ring shows presence of the integrated resistance plasmid and the outer ring denotes community-membership of the population graph shown in (c). Communities match the tree topology, with several basal isolates (n = 11) and a single derived clade ST772-A (n = 329) composed of three population subgroups (A1 – A3). Isolates from two outbreaks in neonatal intensive care units in Ireland are indicated in grey (NICU-1 and NICU-2). Only one representative isolate from longitudinal sampling of a single healthcare worker (n = 39) is included (red circle). **(b)** Basal strains of ST772 showing positions of isolates from India and Bangladesh at the root of the phylogeny (RG28, DAR4066, NKD122). **(c)** Population graph based on pairwise SNP distances, showing SCC*mec* type (node color as for Fig. 1a legend) and population subgroups (polygons, A1-A3). Dashed circles denote hospital-associated outbreaks in Ireland (NICU-1 and NICU-2).

### Emergence and global spread from the Indian subcontinent

Epidemiological and genomic characteristics of ST772 were consistent with an evolutionary origin from the Indian subcontinent. Sixty percent of isolates in this study were collected from patients with family- or travel-background in Bangladesh, India, Nepal or Pakistan, compared to unknown (19%) or other countries (21%) (Fig. 2a, SI Table 2). We found significantly more isolates from India and Bangladesh among the basal strains, compared to clade ST772-A (Fisher’s exact test, 5/11 vs. 47/291, p = 0.026). In particular, three isolates from India and Bangladesh were basal in the (outgroup-rooted) maximum-likelihood phylogeny (Figs. 1b, S1), including two MSSA samples from the original isolations in 2004 (RG28, NKD22). Isolates recovered from South Asia were genetically more diverse than isolates from Australasia and Europe, supporting an origin from the Indian subcontinent (Figs. 2b, S2).

**Fig. 2:**
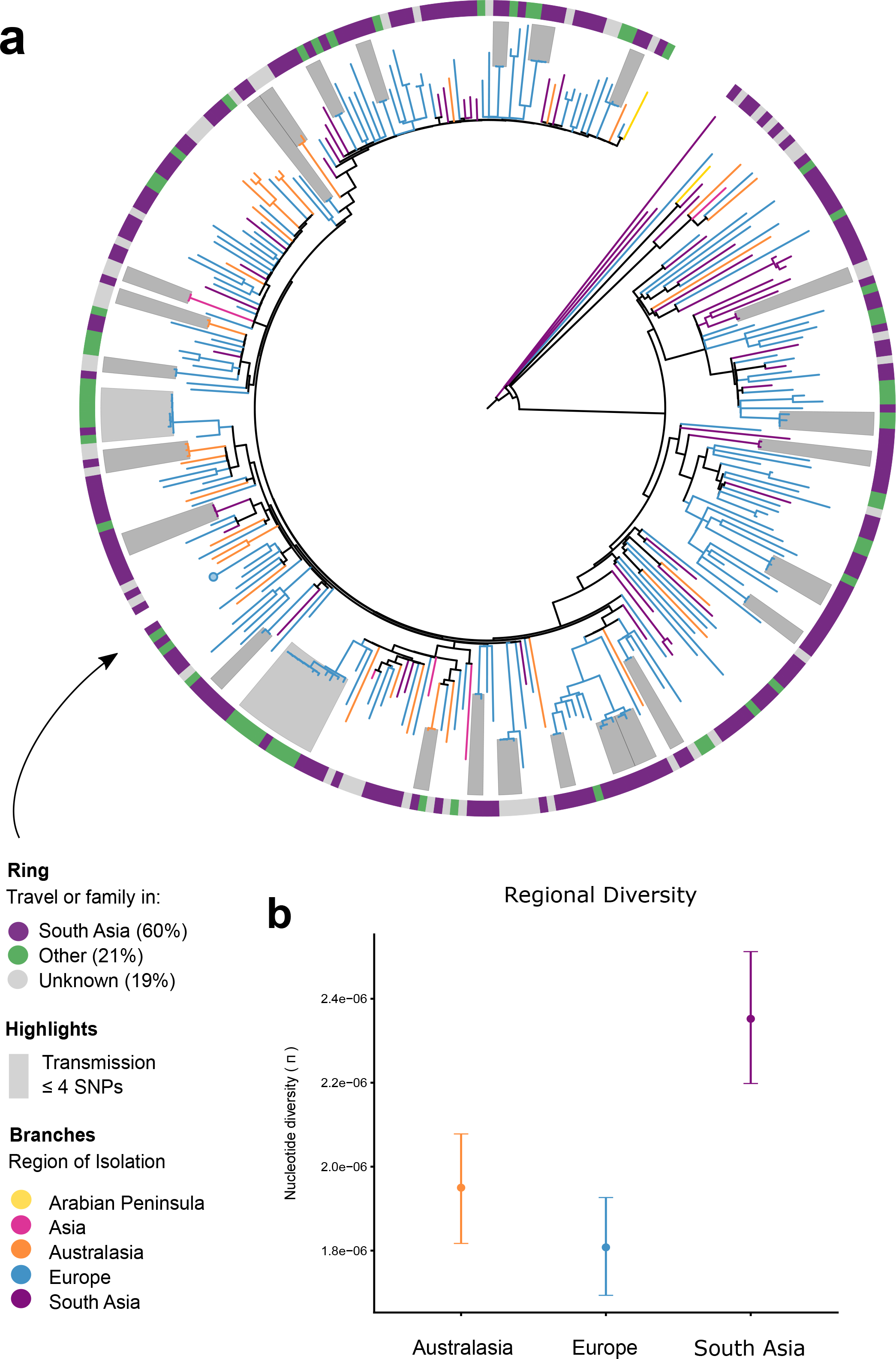
Molecular epidemiology of ST772. (**a**) Patient family- or travel-background in South Asia (India, Pakistan, Nepal, Bangladesh) (59.5%, purple), other countries (21.2%, green) or unknown status (19.3%, gray), is widely distributed across the phylogenetic topology of ST772 (n = 340). Only one representative isolate from longitudinal sampling of a single healthcare worker(n = 39) is included (circle). (**b**) Average pairwise nucleotide diversity per site (π), measured by region (Australasia: orange, n = 36; Europe: blue, n = 244; South Asia: purple, n = 52). Error bars indicate 95% confidence intervals using non-parametric bootstrapping. Isolates from the Arabian Peninsula (n = 2) and Hong Kong (n = 6) were excluded from the diversity analysis due to the small number of samples from these regions.

Consistent with a methicillin-susceptible progenitor, a significantly higher proportion of MSSA was found in the basal isolates (Fisher’s exact test, 4/11 vs. 31/291, p = 0.028) and MSSA isolates demonstrated a lower patristic distance to the root of the maximum likelihood phylogeny compared to MRSA (Fig. S3a). Although it appears that MSSA is proportionately more common in South Asia (Fig. S3b), it is also possible that the observed distribution may be related to non-structured sampling.

Phylogenetic dating suggests an initial divergence of the ancestral ST772 population in 1970 (age of root node: 1970.02, CI: 1955.43 – 1982.60) with a core-genome substitution rate of 1.61 × 10^−6^ substitutions/site/year after removing recombination (Figs. 3a, 3b, S4, S5). This was followed by the emergence of the dominant clade ST772-A and its population subgroups in the early 1990s (ST772-A divergence, 1990.83, 95% CI: 1980.38 – 1995.08). The geographic pattern of dissemination is heterogeneous (Fig. 1a). There was no evidence for widespread endemic dissemination of the clone following intercontinental transmission, although localised healthcare-associated outbreak clusters occurred in neonatal intensive care units in Ireland (NICU-1 and NICU-2, Figs. 1a, S6) (Brennan et al. 2012) and have been reported from other countries in Europe (Blomfeldt et al. 2017) and South Asia (D’Souza et al. 2010; Manoharan et al. 2012; Nadig et al. 2012). While some localised spread in the community was observed among our isolates, patients in local transmission clusters often had traveled to or had family in South Asia (19/27 clusters, Fig. S6).

**Fig. 3:**
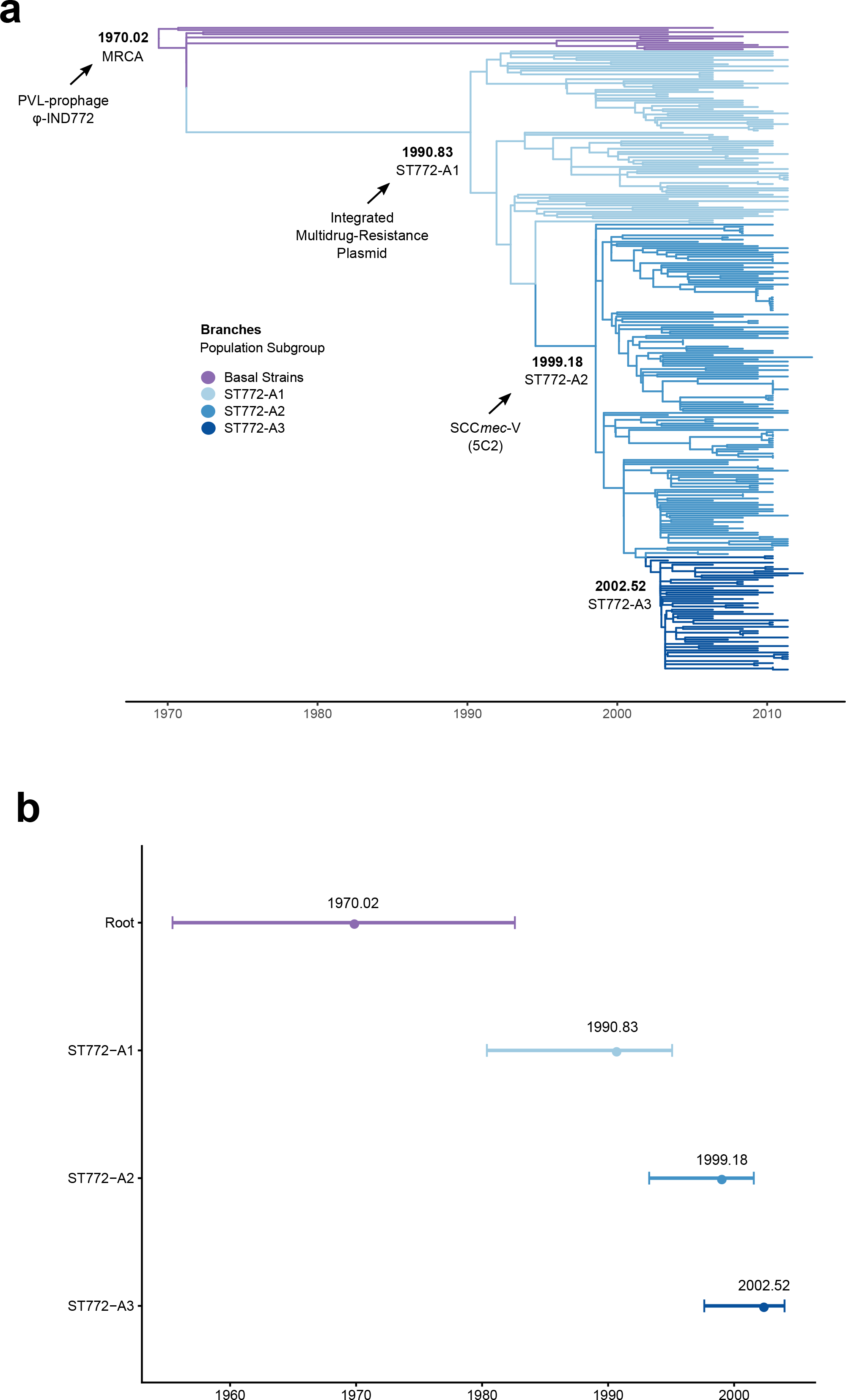
Molecular clock estimates in the emergence of ST772 **(a)** Phylogenetic time-tree with the timescale estimated in Least Squares Dating (LSD). The annotations for nodes represent the time of origin (in years) of basal strains and subgroups A1, A2, A3. Times to the most recent common ancestor (TMRCA) for these lineages are shown. Tips are colored according to the subgroup as per Fig. 1a. The position of the root was optimised during the analysis. Arrows indicate acquisition of three critical mobile genetic elements: the PVL/*sea*-prophage φ-IND772, an integrated multidrug resistance plasmid and the short staphylococcal cassette chromosome SCC*mec*-V (5C2). **(b)** Times to the most recent common ancestor of sub-groups in ST772 after removing recombination. Horizontal bars indicate 95% confidence intervals for nodes (CI) using parametric bootstrapping in LSD.

### Antibiotic resistance acquisition is associated with emergence and dissemination

We examined the distribution of virulence factors, antibiotic resistance determinants and mutations in coding regions to identify the genomic drivers in the emergence and dissemination of ST772. Nearly all isolates (336/340) carried the Panton-Valentine leucocidin (PVL) genes *lukS/F*, most isolates (326/340) carried the associated enterotoxin A (*sea*) and all isolates carried *scn* (SI Table 5). This indicates a nearly universal carriage, across all clades, of both, a truncated *hlb*-converting prophage (the typically associated staphylokinase gene *sak* was only present in one isolate) and the PVL/*sea* prophage φ-IND772 (Prabhakara et al. 2013a). Amongst other virulence factors, the enterotoxin genes *sec* and *sel*, the gamma-hemolysin locus, *egc* cluster enterotoxins and the enterotoxin homologue ORF CM14 were ubiquitous in ST772 (SI Table 7). We detected no statistically significant difference between core virulence factors present in the basal group and ST772-A (SI Table 5, Fig. S7).

We noted a pattern of increasing antimicrobial resistance as successive clades of ST772 emerged. Predicted resistance phenotypes across ST772 were common for ciprofloxacin (97.4%), erythromycin (96.2%), gentamicin (87.7%), methicillin (89.7%), penicillin (100%) and trimethoprim (98.8%), with a corresponding resistome composed of acquired and chromosomally encoded genes and mutations (Figs. 4a, 4b, SI Table 6). There was significantly less predicted resistance in the basal strains compared to ST772-A, including overall multidrug-resistance (≥ 3 classes, 8/11 vs. 291/291, Fisher’s exact test, p < 0.001) (Fig. 5a). The key resistance determinants of interest were the SCC*mec* variants, an integrated resistance plasmid, and other smaller mobile elements and point mutations.

**Fig. 4:**
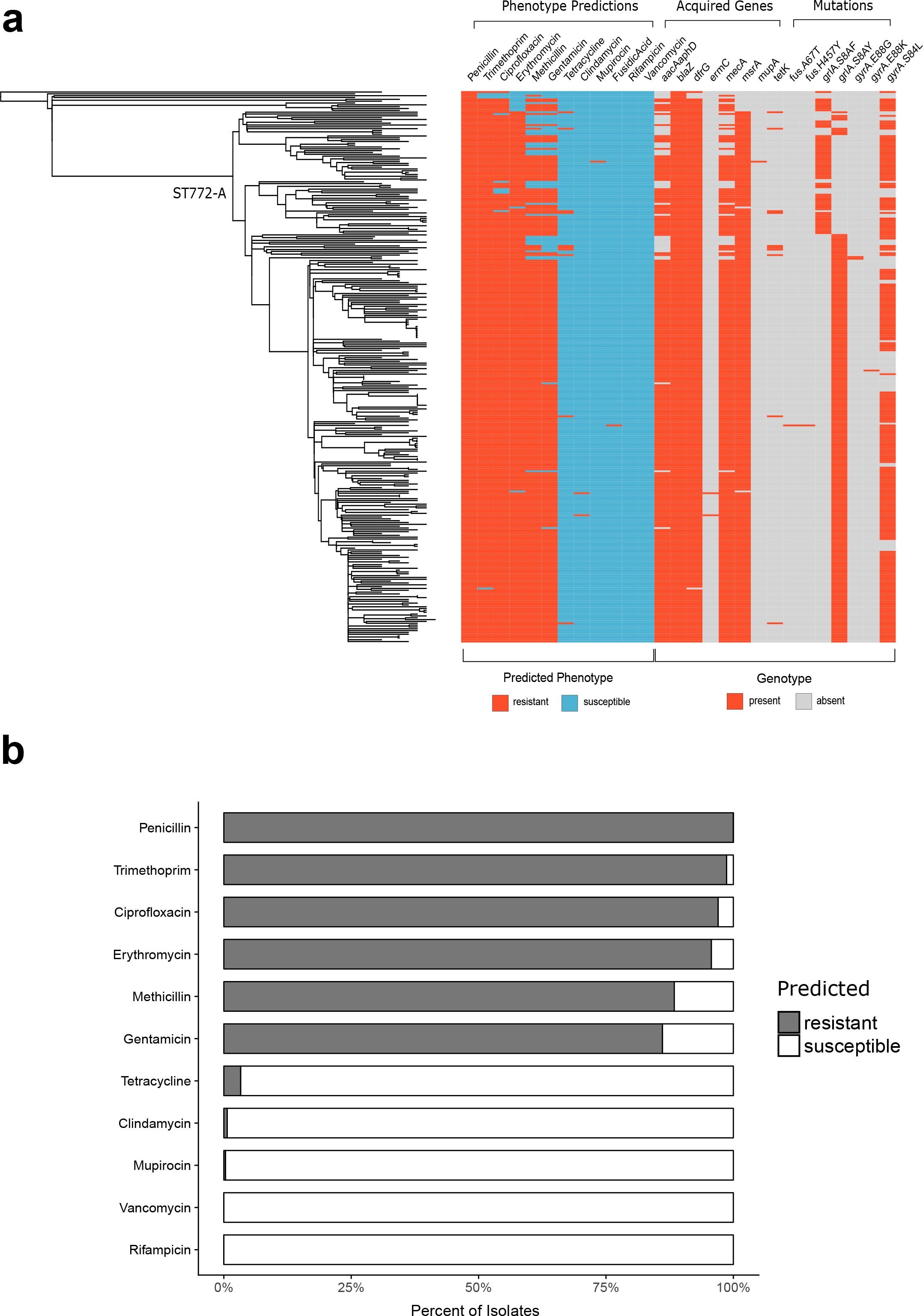
Resistome and predicted resistance phenotypes across ST772. (**a**) Resistome mapped to maximum likelihood phylogeny of ST772. Predicted resistant phenotype is depicted in red, while susceptible phenotype is depicted in blue. Presence of acquired resistance genes and mutations responsible for phenotype predictions are shown in red, while absence of these determinants is shown in gray. (**b**) Percent of isolates predicted resistant (gray) or susceptible (white) for all antimicrobials included in Mykrobe predictor

**Fig. 5:**
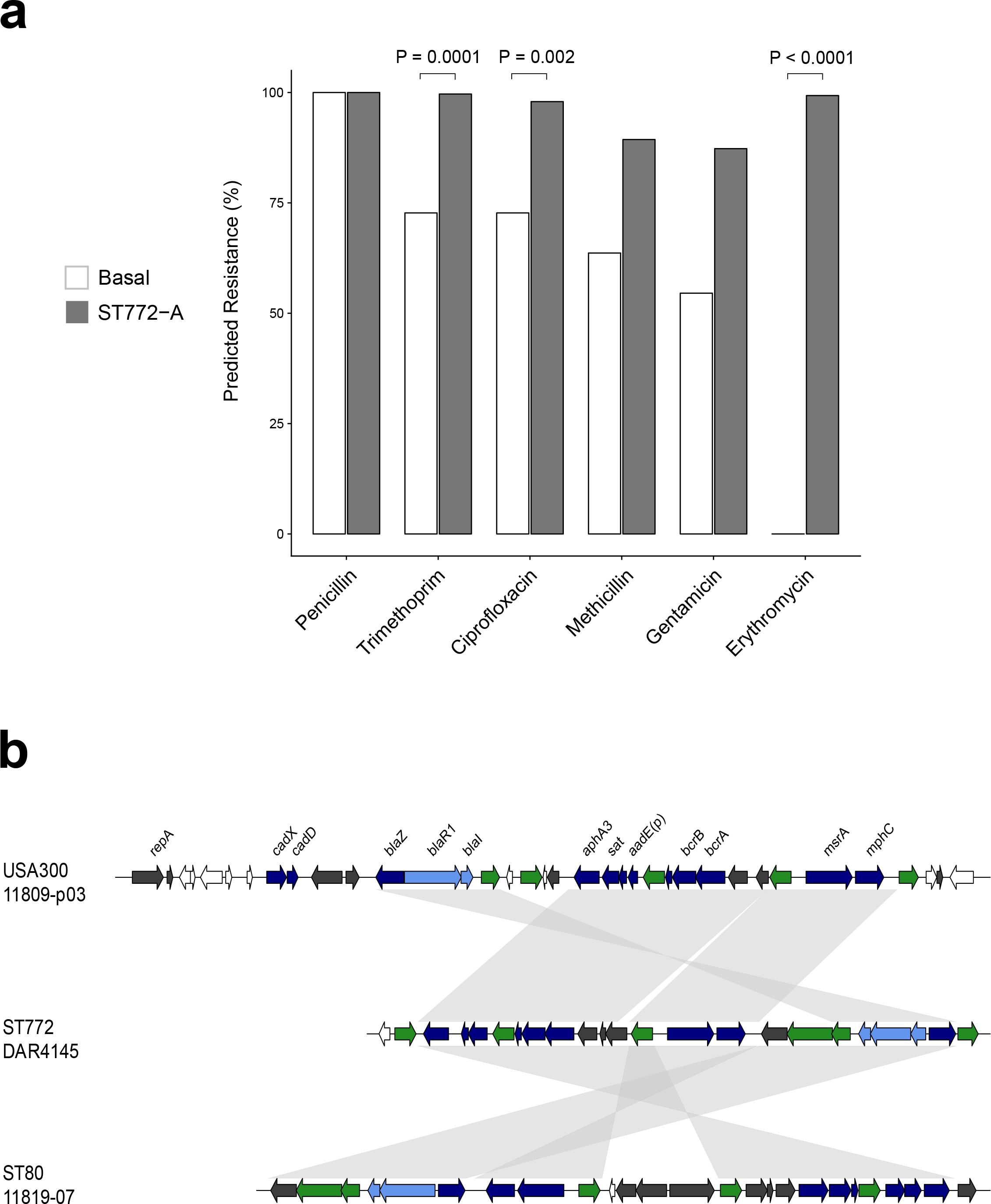
Integrated resistance plasmid in ST772 (**a**) BLAST comparison of the multidrug-resistance plasmid in DAR4145 (middle) with the extrachromosomal plasmid 11809-p03 (top) and the SCC*mec*-IV integrated plasmid in ST80 (bottom), showing alignments > 1000 bp and > 95% nucleotide identity. The comparison highlights three regions harboring resistance genes (dark blue) and their regulators (light blue), which are flanked by transposition elements (green) and appear to have integrated with reversions and rearrangements into ST80 and ST772. Resistance genes include the β-lactam *blaZ* complex, aminoglycoside cluster *aphA3-sat4-aadE* and bacitracin resistance loci *bcrA/B*, as well as macrolide efflux genes *msrA* and *mphC*. Hypothetical proteins and genes of other annotated function are shown in white and dark gray, respectively. **(b)** Proportion of isolates predicted resistant to common antibiotics for basal isolates (white, n = 11) and isolates from ST772-A (gray, n = 291). Values above bars denote statistically significant differences between groups using Fisher’s exact test where p < 0.01.

MRSA isolates predominantly harbored one of two subtypes of SCC*mec*-V: a short variant (5C2) or a composite cassette (5C2&5), which encodes a type 5 *ccr* complex containing *ccrC1* (allele 8) between the *mec* gene complex and *orfX* (Balakuntla et al. 2014) (Fig. S8). Integration of the Tn*4001* transposon encoding aminoglycoside resistance gene *aadA-aphD* occurred across isolates with different SCC*mec* types (260/267), but not in MSSA (0/35). All MRSA isolates (n = 7) within the basal group carried the larger composite cassette SCC*mec*-V (5C2&5), with two of these strains lacking *ccrC* and one isolate carried a remnant of SCC*mec*-IV (Fig. 1a).

The diversity of SCC*mec* types decreased as ST772-A diverged into subgroups (Figs. 1a, 1c, SI Table 6). ST772-A1 included MSSA (n = 30) as well as SCC*mec*-V (5C2) (n = 22) and (5C2&5) (n = 18) strains. Four isolates harbored a putative composite SCC element that included SCC*mec*-V (5C2), as well as *pls* and the *kdp* operon previously known from SCC*mec*-II. One isolate harbored a composite SCC*mec*-V (5C2&5) with copper and zinc resistance elements, known from the European livestock associated CC398-MRSA (Schijffelen et al. 2010). Another six isolates yielded irregular and/or composite SCC elements (SI Table 6). In contrast, the dominant subgroups ST772-A2 and −A3 exclusively carried the short SCC*mec*-V (5C2) element. Eleven of these isolates (including all isolates in NICU-2) lacked *ccrC* and two isolates carried additional recombinase genes (*ccrA/B2* and *ccrA2*).

ST772-A was characterized by the acquisition of an integrated multidrug resistance plasmid (IRP, Fig. 5b), encoding the macrolide-resistance locus *msrA / mphC*, as well as determinants against β-lactams (*blaZ*), aminoglycosides (*aadE-sat4-aphA3*) and bacitracin (*bcrAB*). Thus predicted resistance to erythromycin was uniquely found in ST772-A and not in any of the basal strains (Fisher’s exact test, 289/291 vs 0/11, p < 0.001, Fig. 5a). The mosaic IRP element was highly similar to a composite extrachromosomal plasmid in ST8 (USA300) (Kennedy et al. 2010) and a SCC*mec* integration in the J2 region of the ST80 (Stegger et al. 2012) reference genome (Fig. 5b, SI File 1). A search of closed *S. aureus* genomes (n = 274) showed that the element is rare and predominantly plasmid-associated across ST8 genomes (6/274), with one chromosomal integration in the ST772 reference genome and the SCC*mec* integration in the ST80 reference genome (SI Table 9).

Three basal strains were not multi-drug resistant and included two isolates from the original collections in India (RG28) and Bangladesh (NKD122) (Figs. 1a, 4a). These two strains lacked the trimethoprim determinant *dfrG* and the fluoroquinolone mutations in *grlA* or *gyrA*, encoding only a penicillin-resistance determinant *blaZ* on a Tn*554*-like transposon. However, seven of the strains more closely related to ST772-A did harbor mobile elements and mutations conferring trimethoprim (*dfrG*) and quinolone resistance (*grlA* and *gyrA* mutations). Interestingly, we observed a shift from the quinolone resistance *grlA* S80F mutation in basal strains and ST772-A1, to the *grlA* S80Y mutation in ST772-A2 and -A3 (Fig. 4a).

Thus, the phylogenetic distribution of the key resistance elements suggests acquisition of the IRP by a PVL-positive MSSA strain in the early 1990s (ST772-A1 divergence, 1990.83, 95% CI: 1980.38 – 1995.08), followed by fixation of both the shorter variant of SCC*mec*-V (5C2) and the *grlA* S80Y mutation in a PVL- and IRP-positive MSSA ancestor in the late 1990s (ST772-A2 divergence, 1999.18, 95% CI: 1993.26 – 2001.56) (Figs. 1a, 3a).

### Canonical mutations and phenotypic comparison of basal strains and ST772-A

We found three other mutations of interest that were present exclusively in ST772-A strains (SI Table 7). The first mutation caused a non-synonymous change in *fbpA* (L55P), encoding a fibrinogen-binding protein that mediates surface adhesion in *S. aureus* (Cheung et al. 1995). The second comprised a non-synonymous change (L67V) in the *plc* gene, encoding a phospholipase associated with survival in human blood cells and abscess environments in USA300 (White et al. 2014). The third encoded a non-synonymous mutation (S273G) in *tet(38)*, an efflux pump that promotes resistance to tetracyclines as well as survival in abscess environments and skin colonization (Truong-Bolduc et al. 2014). The functional implication of genes harboring these canonical mutations might suggest a modification of the clone’s ability to colonise and cause SSTIs.

In light of these canonical SNPs, we selected five basal strains and 10 strains from ST772-A to screen for potential phenotypic differences that may contribute to the success of ST772-A. We assessed *in vitro* growth, biofilm formation, cellular toxicity, and lipase activity (Fig. 6, SI Table 8). We found no statistically significant differences between the basal strains and ST772-A in these phenotypic assays, apart from significantly lower lipase activity among ST772-A strains (Welch’s two-sided t-test, t = 3.4441, df = 6.0004, p = 0.0137), which may be related to the canonical non-synonymous mutation in *plc*. However, it is increased rather than decreased lipase activity that has been associated with viability of *S. aureus* USA300 in human blood and neutrophils (White et al. 2014). We found no difference in the median growth rate of ST772-A compared to the basal strains (Mann-Whitney, W = 27, p = 0.8537), although there were two ST772-A strains that grew more slowly suggesting the possibility of some strain to strain variability.

**Fig. 6:**
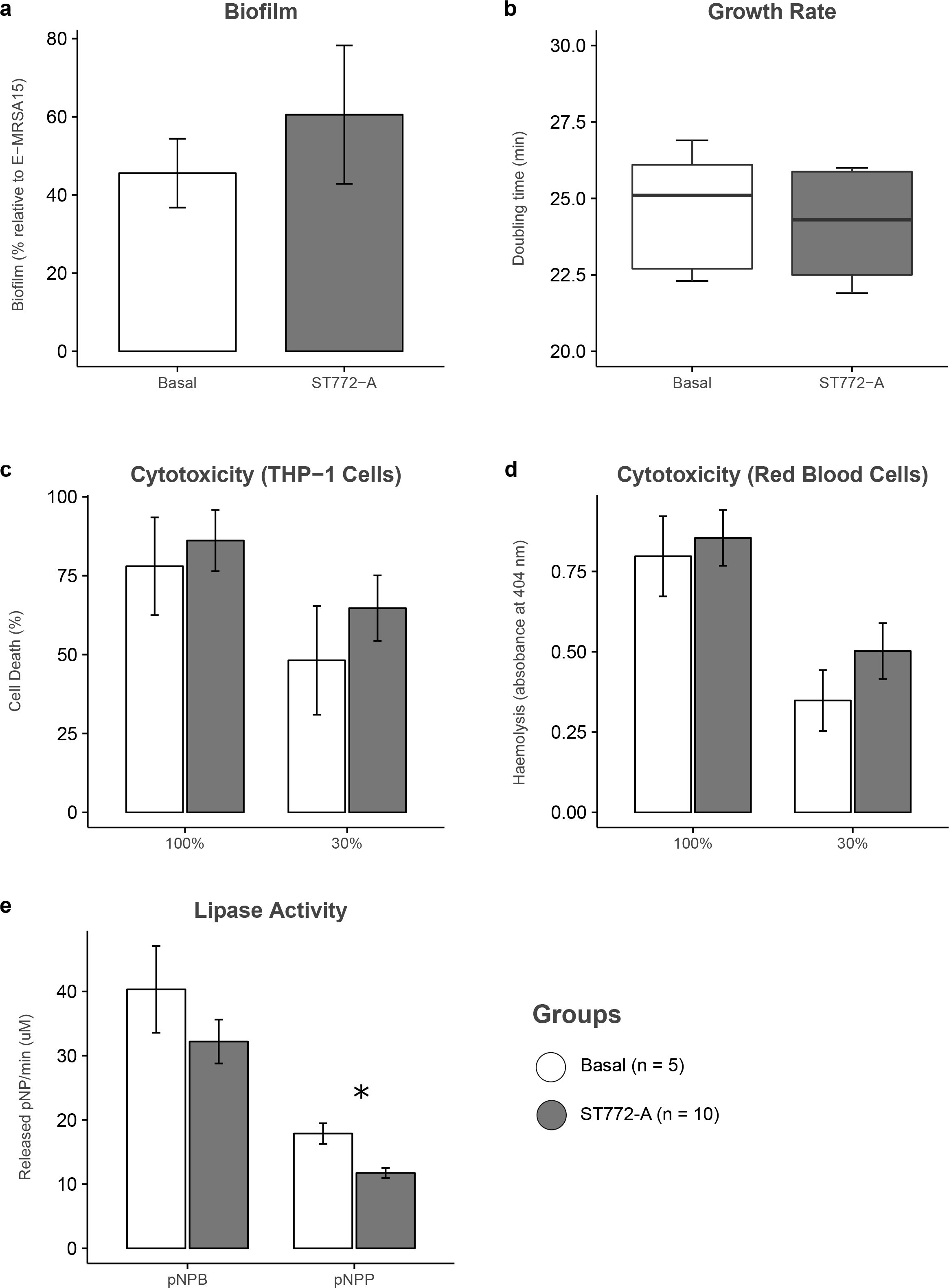
Phenotypic assays for representative strains from the basal group (white, n = 5) and ST772-A (gray, n = 10) for (a) optical density measurements (595 nm) of biofilm formation, accounting for day to day variability relative to control strain E-MRSA15 (%), (b) overnight growth in tryptic soy broth (doubling time per minute) measured by optical density (600 nm), (c) cytotoxicity of neat (100%) and diluted (30%) bacterial supernatant to THP-1 cells measured as cell death by flow cytometry, (d) absorbance measurements (404 nm) of erythrocyte haemolysis in neat (100%) and diluted (30%) bacterial supernatant, (e) lipase activity of *para*-nitrophenyl butyrate (pNPB) or *para*-nitrophenyl palmitate (pNPP) (release of pNP per minute) in neat bacterial supernatant measured by absorbance (410 nm). Slow growing strains H104580604 and HPAS101177P were considered outliers and removed from the growth boxplot for visual clarity after calculation of median and interquartile ranges and assessment of significance. Error bars show standard error; the asterisk denotes a significant difference in pNPP release (Welch’s two-sided t-test, t = 3.4441, df = 6.0004, p = 0.0137) between basal strains and ST772.

## Discussion

In this study, we used whole genome sequencing in combination with epidemiological and phenotypic data to investigate the drivers behind the emergence and spread of a multidrug resistant community-associated MRSA lineage from the Indian subcontinent. Our data suggests that the Bengal Bay clone has acquired the multidrug phenotype of traditional healthcare-associated MRSA, but retains the epidemiological characteristics of community-associated MRSA.

Emergence of a basal population of ST772 appears to have occurred on the Indian subcontinent in the early 1970s and included strains from the original isolations of ST772 in Bangladesh and India in 2004. Recent studies have detected ST772-MSSA and -MRSA in Nepal (Pokhrel et al. 2017) and ST772-MRSA in Pakistan (Madzgalla et al. 2016) also, but it is unclear whether the lineage has been endemic in these countries prior to its emergence in India. Deeper genomic surveillance of ST772-MSSA and –MRSA in the region will be necessary to understand the local epidemiology and evolutionary history of the clone on the Indian subcontinent.

Establishment and expansion of a single dominant clade (ST772-A) occurred in the early 1990s and was associated with the acquisition of an integrated multidrug resistance MGE. The element is similar to a previously described extrachromosomal plasmid of USA300 (Kennedy et al. 2010) and a partially integrated element in the SCC*mec* of a ST80 reference genome (Stegger et al. 2012). While the element was found only once in the ST80 lineage (Stegger et al. 2014) and occurs predominantly on plasmids in closed ST8 (USA300) genomes, its distribution and contribution to the emergence of resistance in the ST8 lineage has so far not been addressed (Planet et al. 2013; Strauß et al. 2017). In contrast, the ubiquitous occurrence and retention of the element in ST772-A suggests that it was instrumental to the emergence of the dominant clade of the Bengal Bay clone.

We observed a lack of significant differences in growth between basal strains and the divergent clade ST772-A. This may suggest that acquisition of drug resistance on this element was not accompanied by a major fitness cost to ST772-A and raises the possibility that members of this clade will survive in environments where antibiotics are heavily used, such as hospitals or in communities with poor antibiotic stewardship, but may also be at little disadvantage in environments where there is less antibiotic use, because its growth rate is comparable to that of non-resistant strains. However, it should be noted that growth assays were conducted *in vitro* and under nutrient-rich conditions that are unlikely to capture anything but very high fitness costs.

Furthermore, we observed a replacement of the long composite SCC*mec*-V (5C2&5) element with the shorter SCC*mec*-V (5C2) and fixation of the quinolone resistance mutation from *grlA* S80F to the *grlA* S80Y as ST772 diverged into its population subgroups in the 1990s. In light of earlier studies demonstrating a fitness advantage in having a smaller SCC*mec* element (Ender et al. 2004; Lee et al. 2007; Collins et al. 2010), the fixation of the shorter SCC*mec*-V (5C2) may be a contributing factor to the success of ST772. We speculate that these changes may have allowed the clone to retain its multidrug resistant phenotype without potentially incurring a significant fitness cost. Further work is required to investigate the role of resistance dynamics in the evolution and fitness potential of ST772.

Given the available epidemiological data, phylogeographic heterogeneity and the clone’s limited success to establish itself in regions outside its endemic range in South Asia (Fig. 1a), there appears to be ongoing exportation of ST772 from the Indian subcontinent, associated with travel and family background in the region. This is supported by reports of MRSA importation in travelers, including direct observations of ST772 importation by returnees from India (Zanger et al. 2012). Our data suggest non-endemic spread within households and the community, including short-term outbreaks at two NICUs in Ireland. This pattern of limited endemic transmission is supported by reports of small transmission clusters in hospitals and households during a comprehensive surveillance study of ST772 in Norway (Blomfeldt et al. 2017). The rapid emergence, global exportation and patterns of local transmission, together with a relatively homogenous genotype, emphasize the clone’s transmissibility.

Overall, the pattern of spread mirrors other community-associated MRSA lineages such as USA300 (Nimmo 2012; Planet et al. 2015), ST80-MRSA (Stegger et al. 2014) and ST59 (Ward et al. 2016) where clones emerge within a particular geographic region, are exported elsewhere, but rarely become established and endemic outside of their place of origin. In contrast, healthcare-associated MRSA clones such as CC22-MRSA-IV (EMRSA-15) (Holden et al. 2013) and ST239-MRSA-III (Harris et al. 2010; Castillo-Ramírez et al. 2012) demonstrate much stronger patterns of phylogeographic structure, consistent with importation into a country followed by local dissemination through the healthcare system. While there are indications for resistance acquisition driving regional community-asscoiated lineages, such as the Taiwan clade of ST59 (Ward et al. 2016), we found strong evidence in our study that the acquisition of mutations and mobile elements associated with multidrug resistance were the dominant driver behind the emergence of the Bengal Bay clone on the Indian subcontinent and its subsequenct intercontinental transmission. Moreover, we observed an unusual, near complete lack of phylogeographic structure in the population, compared to other previously investigated community-associated clones, providing evidence for ongoing circulation and exportation from the Indian subcontinent followed by limited endemic transmission.

Our data traces the evolution of the Bengal Bay clone on the Indian subcontinent, where it emerged in the 1970s and diverged into a single dominant clade in the 1990s, becoming the dominant community-associated MRSA lineage in India. Its rapid emergence was driven by the dissemination of mobile genetic elements, particularly those that confer drug-resistance, such as the acquisition of a multidrug-resistance integrated plasmid and variants of SCC*mec*. Patient epidemiology and phylogenetic heterogeneity suggest a pattern of ongoing exportation from the Indian subcontinent and limited endemic transmission after importation. The Bengal Bay clone therefore appears to combine the epidemiological characteristics of community-associated MRSA lineages with an unusually resistant genotype traditionally seen in healthcare-associated MRSA.

Considering the widespread use of antibiotics and associated poor antibiotic regulation, poor public health infrastructure, and high population density in parts of South Asia, the emergence and global dissemination of multidrug resistant bacterial clones (both Gram-positive and Gram-negative) is alarming, and perhaps not surprising. Global initiatives and funding to monitor the occurrence of emerging clones and resistance mechanisms, and support for initiatives in antimicrobial stewardship at community, healthcare and agricultural levels are urgently needed.

## Materials and Methods

### Isolates

Isolates were obtained from Australia (21), Bangladesh (3), Denmark (70), England (103), Germany (16), Hong Kong (6), India (44), Ireland (28), Italy (2), Netherlands (4), New Zealand (17), Norway (3), Saudi Arabia (1), Scotland (29) and the United Arab Emirates (1) between 2004 and 2012 (SI Table 2). The collection was supplemented with six previously published genome sequences from India (Prabhakara et al. 2012; Prabhakara et al. 2013b; Balakuntla et al. 2014). Notable samples include the initial isolates from Bangladesh and India (Afroz et al. 2008; Goering et al. 2008), two hospital-associated (NICU) clusters from Ireland (Brennan et al. 2012) and longitudinal isolates from a single healthcare worker at a veterinary clinic sampled over two consecutive weeks (VET) (Paterson et al. 2015). Geographic regions were designated as Australasia (Australia, New Zealand), East Asia (Hong Kong), South Asia (India, Bangladesh), Arabian Peninsula (Saudi Arabia, United Arab Emirates) and Europe (Denmark, England, Germany, Ireland, Italy, Netherlands, Norway and Scotland).

### Clinical data and epidemiology

Anonymised patient data was obtained for the date of collection, clinical symptoms, geographic location, epidemiological connections based on family or travel-history, and acquisition in nosocomial- or community-environments, where available (SI Table 2). Clinical symptoms were summarized as SSTI (abscesses, boils, ulcers, exudates, pus, ear and eye infections), urogenital- (vaginal swabs, urine), bloodstream- (bacteremia) or respiratory-infections (pneumonia, lungs abscesses) and colonization (swabs from ear, nose, throat, perineum or environment) (SI Table 2, Fig. S9). Literature and sample maps (SI Maps 1 and 2) were constructed with *geonet*, a wrapper for geographic projections with Leaflet in R (https://github.com/esteinig/geonet).

Where available, acquisition in community- or healthcare-environments was recorded in accordance with guidelines from the CDC. Community-associated MRSA is therein classified as an infection in a person who has none of the following established risk factors for MRSA infection: isolation of MRSA more than 48 h after hospital admission; history of hospitalization, surgery, dialysis or residence in a long-term care facility within one year of the MRSA culture date; the presence of an indwelling catheter or a percutaneous device at the time of culture; or previous isolation of MRSA (Fridkin et al. 2005; Morrison et al. 2006) (Fig. S9).

A valid epidemiological link to South Asia was declared if either travel- or family-background could be reliably traced to Bangladesh, India, Nepal or Pakistan. If both categories (travel and family) were unknown or one did not show a link to the region, we conservatively declared the link as unknown or absent, respectively. The longitudinal collection (n = 39) from a staff member at a veterinary hospital in England was treated as a single patient sample.

### Sequencing, quality control and assembly

Unique index-tagged libraries were created for each isolate, and multiplexed libraries were sequenced on the Illumina HiSeq with 100 bp paired-end reads. Samples from the veterinary staff member were processed and sequenced as described by Paterson et al. (Paterson et al. 2015). Read quality control was conducted with Trimmomatic (Bolger et al. 2014), Kraken (Wood and Salzberg 2014) and FastQC (https://www.bioinformatics.babraham.ac.uk/projects/fastqc). Quality control identified a large proportion of reads classified as *Enterococcus faecalis* in sample HWM2178 (SI Table 3). *In silico* micro-array typing (see below) identified an additional 13 isolates with possible intra-specific contamination due to simultaneous presence of *agr I* and *II*, as well as capsule types 5 and 8 (SI Table 2). We excluded these isolates from all genomic analyses. Raw Illumina data were sub-sampled to 100x coverage and assembled with the SPAdes (Bankevich 2012) pipeline Shovill (https://github.com/tseemann/shovill), which wraps SPAdes, Lighter (Song et al. 2014), FLASH (Magoč and Salzberg 2011), BWA MEM (Li 2013), SAMtools (Li et al. 2009), KMC (Deorowicz et al. 2015) and Pilon (Walker et al. 2014). Final assemblies were annotated with Prokka v1.11(Seemann 2014).

### MLST and SCC typing

*In silico* multi-locus sequence typing (MLST) was conducted using mlst (https://github.com/tseemann/mlst) on the assembled genomes with the *S. aureus* database from PubMLST (https://pubmlst.org/saureus/). Three single locus variants (SLVs) of ST772 were detected and retained for the analysis, describing sequence types ST1573, ST3362 and ST3857 (SI Table 2). Sequences of experimentally verified sets of probes for SCC-related and other *S. aureus* specific markers (Monecke et al. 2011; Monecke et al. 2016) were blasted against SPAdes assemblies (*in silico* micro-array typing), allowing prediction of presence or absence of these markers and detailed typing of SCC elements. We assigned MRSA to four isolates that failed precise SCC classification based on presence of *mecA* on the probe array and detection of the gene with Mykrobe predictor (Bradley et al. 2015).

### Variant calling

Samples passing quality control (n = 340) were aligned to the PacBio reference genome DAR4145 from Mumbai and variants were called with the pipeline Snippy (available at https://github.com/tseemann/snippy) which wraps BWA MEM, SAMtools, SnpEff (Cingolani et al. 2012) and Freebayes (Garrison and Marth 2012). Core SNPs were defined as being present in all samples (ignoring insertions and deletions, n = 7,063) and were extracted with *snippy-core* at default settings. We assigned canonical SNPs for ST772-A, as those present exclusively in all isolates of ST772-A, but not in the basal strains. Annotations of variants were based on the reference genome DAR4145.

### Phylogenetics and recombination

A maximum-likelihood (ML) tree under the General Time Reversible model of nucleotide substitution with among-site rate heterogeneity across 4 categories (GTR + Γ), ascertainment bias correction (Lewis) and 100 bootstrap (BS) replicates was generated based on 7,063 variant sites (core-genome SNPs) in RaxML-NG 0.5.0 (available at https://github.com/amkozlov/raxml-ng), which implements the core functionality of RAxML (Stamatakis 2014). The tree with the highest likelihood out of ten replicates was midpoint-rooted and visualized with interactive Tree of Life (ITOL) (Figs. 1a, 2a, S6, S12a) (Letunic and Bork 2007). In all phylogenies (Figs. 1a, 2a, 3a, S6, S10, S12a) samples from the veterinary staff member were collapsed for clarity.

A confirmation alignment (n = 351) was computed as described above for resolving the pattern of divergence in the basal strains of ST772. The alignment included the CC1 strain MW2 as outgroup, as well as another known SLV of CC1, sequence type 573 (n = 10). The resulting subset of core SNPs (n = 25,701) was used to construct a ML phylogeny with RaxML-NG (GTR + Γ) and 100 bootstrap replicates (Fig. S1). We also confirmed the general topology of our main phylogeny as described above using the whole genome alignment of 2,545,215 nucleotides generated by Snippy, masking sites if they contained missing (−) or uncertain (N) characters across ST772.

Gubbins (Croucher et al. 2015) was run on a complete reference alignment with all variant sites defined by Snippy to detect homologous recombination events, using a maximum of five iterations and the GTR + Γ model in RaxML (Fig. S10). A total of 205 segments were identified as recombinant producing a core alignment of 7,928 SNPs. Phylogenies were visualized using ITOL, *ape* (Paradis et al. 2004), *phytools* (Revell 2012), *ggtree* (Yu et al. 2017) or *plotTree* (https://github.com/holtlab/plotTree/). Patristic distances to the root of the phylogeny (Fig. S2) were computed in the *adephylo* (Jombart et al. 2010) function *distRoot*.

### Dating analysis

We used LSD v0.3 (To et al. 2016) to obtain a time-scaled phylogenetic tree. This method fits a strict molecular clock to the data using a least-squares approach. Importantly, LSD does not explicitly model rate variation among lineages and it does not directly account for phylogenetic uncertainty. However, its accuracy is similar to that obtained using more sophisticated Bayesian approaches (Duchêne et al. 2016), with the advantage of being computationally less demanding.

LSD typically requires a phylogenetic tree with branch lengths in substitutions per site, and calibrating information for internal nodes or for the tips of the tree. We used the phylogenetic tree inferred using Maximum likelihood in PhyML (Guindon et al. 2010) (before and after removing recombination with Gubbins, as described above) using the GTR+Γ substitution model with 4 categories for the Γ distribution. We used a combination of nearest-neighbour interchange and subtree-prune-regraft to search tree space. Because PhyML uses a stochastic algorithm, we repeated the analyses 10 times and selected that with the highest phylogenetic likelihood. To calibrate the molecular clock in LSD, we used the collection dates of the samples (i.e. heterochronous data). The position of the root can be specified *a priori*, using an outgroup or by optimising over all branches. We chose the latter approach. To obtain uncertainty around node ages and evolutionary rates we used the parametric bootstrap approach with 100 replicates implemented in LSD.

An important aspect of analysing heterochronous data is that the reliability of estimates of evolutionary rates and timescales is contingent on whether the data have temporal structure. In particular, a sufficient amount of genetic change should have accumulated over the sampling time. We investigated the temporal structure of the data by conducting a regression of the root-to-tip distances of the Maximum likelihood tree as a function of sampling time (Korber et al. 2000), and a date-randomisation test (Ramsden et al. 2009). Under the regression method, the slope of the line is a crude estimate of the evolutionary rate, and the extent to which the points deviate from the regression line determines the degree of clocklike behaviour, typically measured using the R (Rambaut et al. 2016). The date randomisation test consists in randomising the sampling times of the sequences and re-estimating the rate each time. The randomisations correspond to the distribution of rate estimates under no temporal structure. As such, the data have strong temporal structure if the rate estimate using the correct sampling times is not within the range of those obtained from the randomisations (Duchêne et al. 2015). We conducted 100 randomisations, which suggested strong temporal structure for our data (Fig. S3). We also verified that the data did not display phylogenetic-temporal clustering, a pattern which sometimes misleads the date-randomisation test (Murray et al. 2016).

Results from this analysis (substitution rates, and node age estimates) using phylogenies before and after removing recombination were nearly identical (Figs. S4, S5). We therefore chose to present results from our analysis after removing recombination.

### Nucleotide diversity

Pairwise nucleotide diversity and SNP distance distributions for each region with n > 10 (Australasia, Europe, South Asia) were calculated as outlined by Stucki et al. (Stucki et al. 2016). Pairwise SNP distances were computed using the SNP alignment from Snippy (n = 7,063) and the *dist.dna* function from *ape* with raw counts and deletion of missing sites in a pairwise fashion. An estimate of average pairwise nucleotide diversity per site (π) within each geographic region was calculated from the SNP alignments using raw counts divided by the alignment length. Confidence intervals for each region were estimated using 1000 bootstrap replicates across nucleotide sites in the original alignment via the *sample* function (with replacement) and 2.5% - 97.5% quantile range (Fig. 2b).

### Population structure

We used the network-analysis and -visualization tool NetView (Neuditschko et al. 2012; Steinig et al. 2016) (available at http://github.com/esteinig/netview) to delineate population subgroups in ST772. Pairwise Hamming distances were computed from the core SNP alignment derived from Snippy. The distance matrix was used to construct mutual k-nearest-neighbour networks from *k* = 1 to *k* = 100. We ran three commonly used community detection algorithms as implemented in *igraph* to limit the parameter choice to an appropriate range for detecting fine-scale population structure: fast-greedy modularity optimization (Girvan and Newman 2002), Infomap (Rosvall and Bergstrom 2008) and Walktrap (Pons and Latapy 2006). We thereby accounted for differences in the mode of operation and resolution of algorithms. Plotting the number of detected communities against *k*, we were able to select a parameter value at which the results from the community detection were approximately congruent (Fig. S11).

Since we were interested in the large-scale population structure of ST772, we selected *k* = 40 and used the low-resolution fast-greedy modularity optimisation to delineate final population subgroups. Community assignments were mapped back to the ML phylogeny of ST772 (Fig. 1a). All subgroups agreed with the phylogenetic tree structure and were supported by ≥ 99% bootstrap values (Fig. S12). One exception was isolate HW_M2760 located within ST772-A2 by phylogenetic analysis, but assigned to ST772-A3 by network analysis (Figs. S11, S12). This appeared to be an artefact of the algorithm, as its location and connectivity in the network representation matched its phylogenetic position within ST772-A2. The network and communities were visualized using the Fruchtermann-Reingold algorithm (Fig. 1c), excluding samples from the veterinary staff member in Fig. 1c (Fig. S11).

### Local transmission clusters

We obtained approximate transmission clusters by employing a network approach supplemented with the ML topology and patient data, including date of collection, location of collection and patient family links and travel or family links to South Asia. We used pairwise SNP distances to define a threshold of 4 SNPs, corresponding to the maximum possible SNP distance obtained within one year under a core genome substitution rate of 1.61 × 10^−6^ nucleotide substitutions/site/year. We then constructed the adjacency matrix for a graph, in which isolates were connected by an undirected edge, if they had a distance of less or equal to 4 SNPs. All other isolates were removed from the graph and we mapped the resulting connected components to the ML phylogeny, showing that in each case the clusters were also reconstructed in the phylogeny, where isolates diverged from a recent common ancestor (gray highlights, Figs. 2a, S6). We then traced the identity of the connected components in the patient meta-data and added this information to each cluster. NICU clusters were reconstructed under these conditions (Figs. 2a, S6).

### Antimicrobial resistance, virulence factors and pan-genome analysis

Mykrobe Predictor was employed for antibiotic susceptibility prediction and detection of associated resistance determinants and mutations. Mykrobe Predictor has demonstrated sensitivity and specificity > 99% for calling phenotypic resistance and is comparable to gold-standard phenotyping in *S. aureus* (Bradley et al. 2015). Predicted phenotypes were therefore taken as a strong indication for actual resistance phenotypes in ST772. Genotype predictions also reflect multidrug resistance profiles (aminoglycosides, β-lactams, fluoroquinolones, MLS, trimethoprim) reported for this clone in the literature (Ellington et al. 2010; Brennan et al. 2012; Chakrakodi et al. 2014; Shore et al. 2014; Steinig et al. 2015; Blomfeldt et al. 2017). As most resistance-associated MGEs in the complete reference genome DAR4145 are mosaic-like and flanked by repetitive elements (Steinig et al. 2015), we used specific diagnostic genes present as complete single copies in the reference annotation of DAR4145 (Steinig et al. 2015) to define presence of the IRP (*msrA*) and Tn*4001* (*aacA-aphD*). Mykrobe Predictor simultaneously called the *grlA* mutations S80F and S80Y for quinolone resistant phenotypes. However, in all cases one of the variants was covered at extremely low median k-mer depth (< 20) and we consequently assigned the variant with higher median k-mer depth at *grlA* (SI Table 6).

ARIBA (Hunt et al. 2017) with default settings and the core Virulence Factor database were used to detect the complement of virulence factors in ST772. We corroborated and extended our results with detailed *in-silico* microarray typing, including the presence of the *egc* gene cluster or *S. aureus* specific virulence factors such as the enterotoxin homologue ORF CM14. Differences in detection of relevant virulence factors between the *in silico* typing and ARIBA included, amongst others, *lukS/F-PVL* (337 vs. 336), *sea* carried on the φ-IND772 prophage (336 vs. 326), *sec* (333 vs 328) and *sak* (1 vs. 2). Since *in silico* microarray typing was based on assembled genomes and may therefore be prone to assembly errors, we used results from the read-based typing with ARIBA to assess statistical significance of virulence factors present in basal strains and ST772-A (Fig. S7).

Pan-genome analysis was conducted using Prokka annotated assemblies in Roary (Page et al. 2015), with minimum protein BLAST identity at 95% and minimum percentage for a gene to be considered core at 99% (Fig. S13). A gene synteny comparison between major SCC*mec* types was plotted with genoPlotR(Guy et al. 2010) (Fig. S8). A nucleotide BLAST comparison between the extrachromosomal plasmid 11809-03 of USA300, the integrated resistance plasmid in the ST772 reference genome DAR4145 and the integrated plasmid region in strain 11819-07 of ST80 was plotted with geneD3 (https://github.com/esteinig/geneD3/), showing segments > 1kb (SI File 1).

We searched for the three resistance regions which aligned to the 11819-07 and the 11809-03 plasmid (DAR4145 reference genome; R1: 1456024-1459959 bp, R2: 1443096-1448589 bp and R3: 1449679-1453291 bp) in all completed *S. aureus* genomes (including plasmids) in RefSeq (NCBI) and the NCTC3000 project (http://www.sanger.ac.uk/resources/downloads/bacteria/nctc/) using nctc-tools (https://github.com/esteinig/nctc-tools) and nucleotide BLAST with a minimum of 90% coverage and identity (n = 273). Since the IRP is mosaic-like and composed of several mobile regions, we only retained query results, if all three of the regions were detected (SI Table 9). We then traced the integration sites in the accessions, determining whether integrations occurred the chromosome or plasmids. Multi-locus sequence types were assigned using *mlst* (https://github.com/tseemann/mlst).

### Growth curves

S. *aureus* strains were grown overnight in 5 mL tryptic soy broth (TSB, Fluka) with shaking (180 *rpm*) at 37 °C. Overnight cultures were diluted 1:1000 in fresh TSB and 200 μL was added to a 96 – well plate (Costar) in triplicate. Growth was measured 37 °C, with shaking (300 *rpm*) using a FLUOROstar fluorimeter (BMG Labtech) using an absorbance wavelength of 600 nm. Growth curves represent the mean of triplicate results.

### Cell culture conditions

The monocyte-macrophage THP-1 cell line was maintained in suspension in 30 mL Roswell Park Memorial Medium Institute (RPMI-1640) medium, supplemented with 10% heat-inactivated fetal bovine serum (FBS), 1 μM L-glutamine, 200 units/mL penicillin, and 0.1 mg/mL streptomycin at 37 °C in a humidified incubator with 5% CO_2_. Cells were harvested by centrifugation at 700 × *g* for 10 min at room temperature and re-suspended to a final density of 1−1.2 × 10^6^ cells/mL in tissue-grade phosphate buffered saline, typically yielding >95 % viable cells as determined by easyCyte flow cytometry (Millipore).

Human erythrocytes were harvested from 10 mL of human blood following treatment in sodium heparin tubes (BD). Whole blood was centrifuged at 500 × *g* for 10 min at 4 °C. Supernatant (plasma) was aspirated and cells were washed twice in 0.9 % NaCl and centrifuged at 700 × *g* for 10 min. Cell pellet was gently re-suspended in 0.9 % NaCl and diluted to 1 % (v/v).

### Cytotoxicity assay

To monitor *S. aureus* toxicity, *S. aureus* strains were grown overnight in TSB, diluted 1:1000 in 5 mL fresh TSB and grown for 18 h at 37 °C with shaking (180 *rpm*). Bacterial supernatants were prepared by centrifugation of 1 mL of bacterial culture at 20,000 × *g* for 10 min. For assessing toxicity to THP-1 cells, 20 μL of cells were incubated with 20 μL of bacterial supernatant and incubated for 12 min at 37 °C. Both neat and 30% diluted supernatant (in TSB) were used as certain *S. aureus* strains were considerably more toxic than others. Cell death was quantified using easyCyte flow cytometry using the Guava viability stain according to manufacturer’s instructions. Experiments were done in triplicate. For assessing haemolysis, 150 μL of 1% (v/v) erythrocytes were incubated with 50 μl of either neat and 30% supernatant in a 96 well plate for 30 min at 37°C. Plates were centrifuged for 5 min at 300 × *g* and 75 μL of supernatant was transferred to a new plate and absorbance was measured at 404nm using a FLUOROstar fluorimeter (BMG Labtech). Normalised fluorescence was achieved using the equation (A_t_−A_0_) / (A_m_ / A_0_) where A_t_ is the haemolysis absorbance value of a strain, A_0_ is the minimum absorbance value (negative control of 0.9% NaCl) and Am is the maximum absorbance value (positive control of 1 % triton X-100).

### Lipase assay

Bacterial supernatants used in the above cytotoxicity assays were also used to assess lipase activity, using the protocol published by Cadieux *et al.* (Cadieux et al. 2014) with modifications. Briefly, 8mM *para*-nitrophenyl butyrate (pNPB), the short chain substrate, or *para*-nitrophenyl palmitate (pNPP), the long chain substrate, (Sigma) was mixed with a buffer (50mM Tris-HCl (pH 8.0), 1mg/ml gum Arabic and 0.005% Triton-X100) in a 1:9 ratio to create assay mixes. A standard curve using these assay mixes and *para*-nitrophenyl (pNP) (Sigma) was created, and 200μl of each dilution was pipetted into one well of a 96-well plate (Costar). 180μl of the assay mixes was pipetted into the remaining wells of a 96-well plate, and 20μl of the harvested bacterial supernatant was mixed into the wells. The plate was placed in a FLUOstar Omega microplate reader (BMG Labtech) at 37°C, and a reading at 410nm was taken every 5 min.s for 1h. The absorbance readings were converted to μM pNP released/min. using the standard curve.

### Biofilm formation

Semi-quantitative measurements of biofilm formation on 96-well, round-bottom, polystyrene plates (Costar) was determined based on the classical, crystal violet method of Ziebuhr et al. (Ziebuhr et al. 1997). 18 h bacterial cultures grown in TSB were diluted 1:40 into 100 μL TSB containing 0.5 % glucose. Perimeter wells of the 96-well plate were filled with sterile H2O and plates were placed in a separate plastic container inside a 37°C incubator and grown for 24 h under static conditions. Following 24 h growth, plates were washed five times in PBS, dried and stained with 150 μL of 1% crystal violet for 30 min at room temperature. Following five washes of PBS, wells were re-suspended in 200 μL of 7% acetic acid, and optical density at 595 nm was measured using a FLUOROstar fluorimeter (BMG Labtech). To control for day to day variability, a control strain (E-MRSA15) was included on each plate in triplicate, and absorbance values were normalised against this. Experiments were done using six technical repeats from 2 different experiments.

### Statistical analysis

All statistical analyses were carried out in R or python and considered significant at p < 0.05, except for comparisons of proportions across the multiple virulence and resistance elements, which we considered be statistically significant at p < 0.01. Veterinary samples (n = 39) were restricted to one isolate (one patient, Staff_E1A) for statistical comparison of region of isolation, proportion of resistance, virulence and MSSA between basal strains and ST772-A (n = 302, main text, Figs. 5a, S7). Differences in pairwise SNP distance and nucleotide diversity between all regions were assessed using non-parametric Kruskal-Wallace test and post-hoc Dunn’s test for multiple comparisons with Bonferroni correction, as distributions were assumed to be not normally distributed (Fig. 2b, n = 340, Fig. S2). Phenotypic differences were assessed for normality with Shapiro-Wilk tests. We consequently used either Welch’s two-sided t-test or the non-parametric two-sided Wilcoxon rank-sum test (Fig. 6, SI Table 8).

## Data availability

Core analyses, including parameter settings, cluster resource configurations and versioned software distributions are reproducible through the *bengal-bay-0.1* workflow, which can be found along with other scripts and data files at our GitHub repository (https://github.com/esteinig/ST772). The workflow is implemented in Snakemake (Köster and Rahmann 2012) and runs in virtual environments that include software distributed in the Bioconda (Dale et al. 2017) channel. Workflow results are summarized in the Supplementary Tables. Analyses were conducted on the Cheetah cluster at Menzies School of Health Research, Darwin.

Short-read sequences have been deposited at ENA under accession numbers detailed in SI Table 2. Additional isolates from India are available from the SRA under accession numbers SRR404118, SRR653209, SRR653212 and SRR747869-SRR747873. Outgroup strains used in the context phylogeny are available from ENA under accession numbers SRR592258 (MW2), ERR217298, ERR217349, ERR221806, ERR266712, ERR279022, ERR279023, ERR278908. ERR279026, ERR716976, ERR717011 (ST573). The ST772 reference genome DAR4145 is available at GenBank under accession number CP010526.1.

## Author contributions

EJS, ST conducted the bioinformatics analysis; SD performed the dating analysis; MY, ML, RM conducted phenotyping experiments; SM, PS, PA performed *in silico* typing and provided bioinformatics support; DS provided support on the computing cluster; DAR, DW, AK, RG, ED, RE, SM, MI, MO, GC, AP, GB, AS, DC, AP, AM, HdL, HW, NK, HH, BS, FL, SP, SW, HA, LS, SH provided strains and relevant meta-data; EJS, ST, DAR, SM, MTGH wrote the manuscript; all authors contributed to critical review of the manuscript. ST directed the project with support from SB and JP.

## Acknowledgements

We thank the library construction, sequencing, and core informatics teams at the Wellcome Trust Sanger Institute. We also extend our gratitude to Anand Manoharan for comments on the manuscript and strains from India. This work was supported by the Australian National Health and Medical Research Council (#1145033 to ST, #1065908 to MO), by the National Institute of Health grant (GM080602 to DAR), the Irish Health Research Board (HRA-POR-2015-1051 to DC and AS).

