## Supplementary Figures for "Evolution and global transmission of a multidrug-resistant, community-associated MRSA lineage from the Indian subcontinent"

**Supplementary Maps and Files**

All supplementary files can be found at: https://doi.org/10.6084/m9.figshare.8061887.v1

Please note that you will need an active internet connection to view the interactive files. Maps and the interactive gene comparison can be viewed in all modern browsers (Firefox, Chrome, Edge, Opera).

**Map 1**: Interactive map of publications on ST772 by country (circles, coloured by region; blue: Europe, seagreen: Australasia, purple: Southeast Asia, violet: South Asia, green: Americas, orange: Middle East, red: Africa) generated with Leaflet for R. Circles can be clicked to show publication information and link to PubMed. Circle coordinates were randomly jittered by 0.01° longitude / latitude for visibility and mapped using non-specific coordinates for the respective countries. A full account of the publications is available in Supplementary Tables 1

**Map 2:** Interactive map of ST772 samples from this study, with country- or city-resolution, where available, (coloured by type; blue: MSSA, red: MRSA) generated with Leaflet for R. Circles can be clicked for information on samples and links to accessions in the SRA. Circle coordinates were randomly jittered by 0.01° longitude / latitude for visibility. A full account of sample meta data is available in Supplementary Tables 2.

**File 1:** Interactive gene comparison of the integrated resistance plasmid region in DAR4145 (middle), the extrachromosomal 18809-p03 in USA300 (top) and the integrated plasmid in the dominant European lineage ST80 (bottom). Segments (predicted coding regions) and nucleotide BLAST comparisons (> 80% identity, > 1kb alignment length) can be clicked for annotations. Transposition regions (green) flank three major regions of the integrated plasmid, which harbours aminoglycoside, beta-lactam and macrolide resistance genes (blue). Resistance regions appear to have inserted into the chromosome of DAR4145 with inversions and rearrangements compared to 18809-p03 and ST80.

**File 2:** High resolution image of epidemiological data, transmission clusters (connected components) and phylogeny of ST772 as outlined in Fig. S6 with accompanying Table. Rings in the phylogeny denote patient family or travel links to South Asia, while node colour of connected components denotes generalised epidemiological link to South Asia (see Methods).

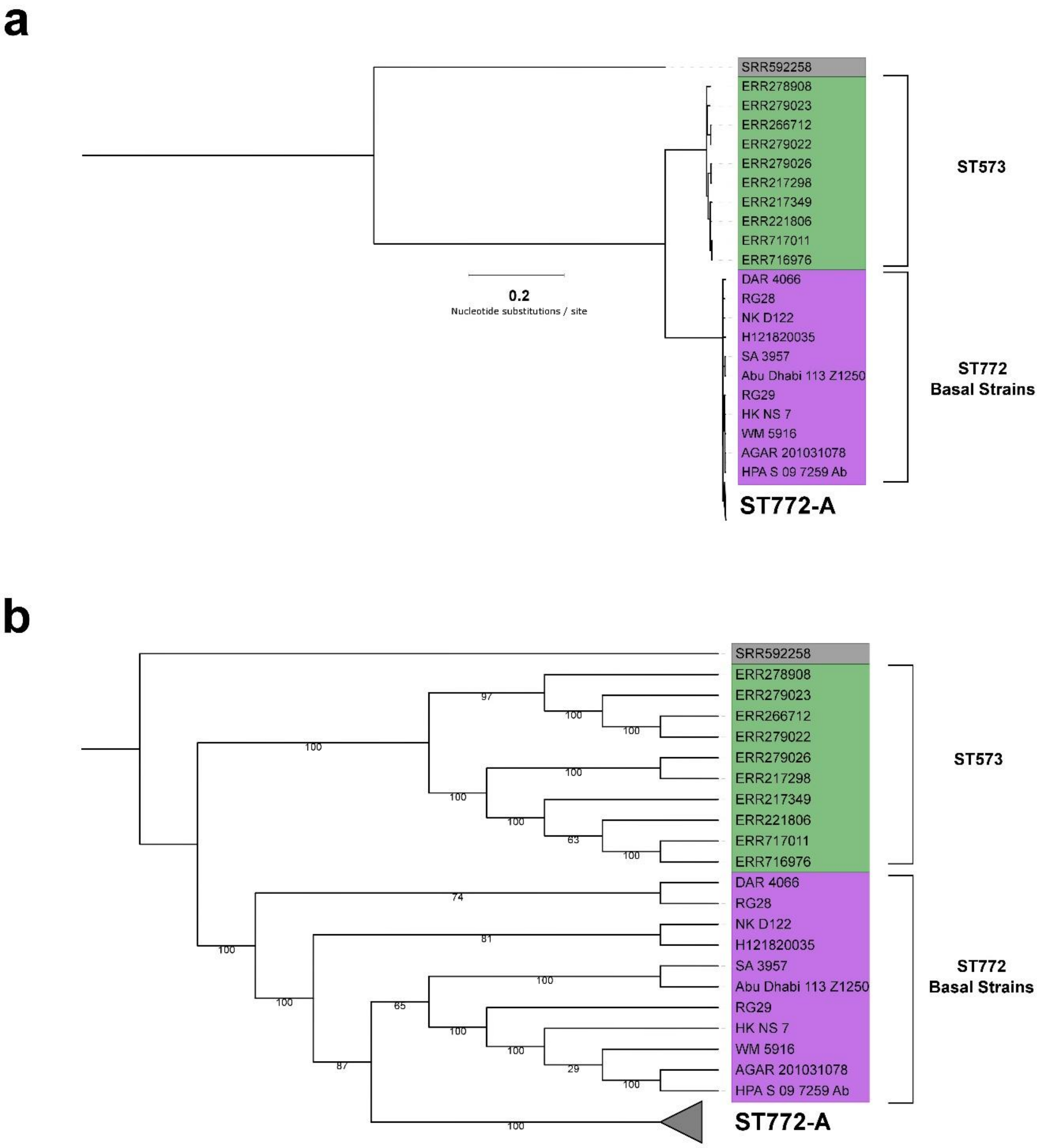


**Fig. S1:** Maximum likelihood (GTR + G) phylogeny based on 25,701 SNPs rooted on outgroup strain MW2 (CC1, SRR592258) and including additional strains from another single locus variant of CC 1, ST573 (n = 10) with branch lengths (a) and as cladogram with 100 bootstrap support values (b). The phylogeny resolves the basal strain topology and is highly similar to the Least Squares Dating estimated root (Fig. S4) and the midpoint-rooted within-lineage phylogeny of ST772 (Figure 1a, 1b) with strains from India at the base of ST772 (DAR4066, RG28, bootstrap support for basal position outside remainder of strains 100%).

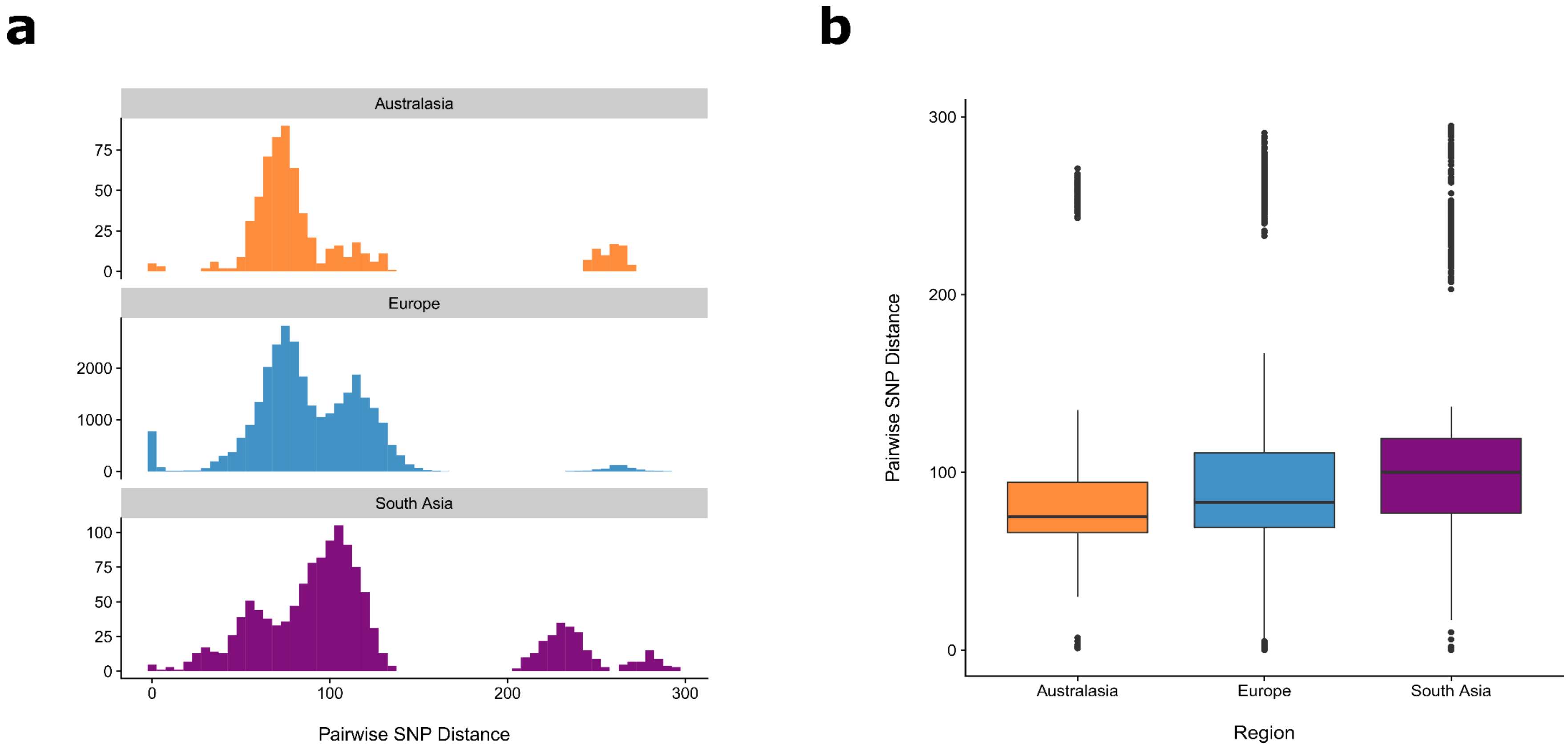


**Fig. S2:** Distribution of pairwise SNP distances between isolates from regions where n > 10 (orange: Australasia, blue: Europe, purple: South Asia) showing (a) histogram representation with a distinction between the dominant clade ST772-A (distribution on the left) and basal strains (distribution on the right) and (b) boxplot of pairwise SNP distance for each region. Kruskal-Wallace test on pairwise SNP distances suggests significant differences between the regions ($\chi^2$= 171.22, df = 2, $p < 1 \times 10^{-6}$). Post-hoc Dunn's test with Bonferroni correction for pairwise multiple comparisons demonstrates significant differences between all pairwise combinations ($p < 1 \times 10^{-6}$).

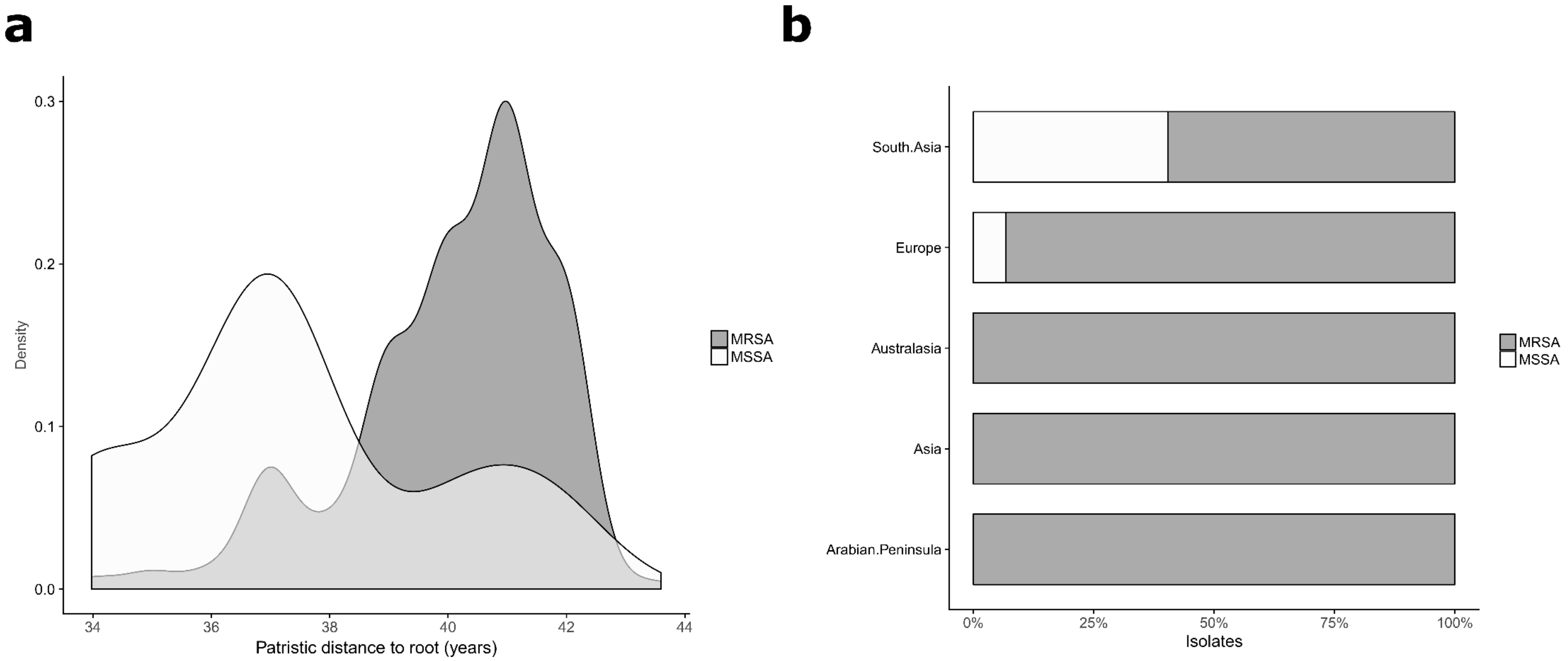


**Fig. S3:** Distribution of methicillin-susceptible ST772 (MSSA). (a) Density distribution of patristic distance (in years using Least Squares Dating (LSD) rooted Maximum Likelihood phylogeny) to the root of the phylogeny for MSSA and MRSA, demonstrating basal positions of MSSA isolates in the LSD-rooted and dated phylogeny of ST772 (Fig. S4a). (b) Proportion of MSSA compared to MRSA isolates within regions. South Asia (n = 52) shows the highest proportion of MSSA isolates in concordance with a hypothesised MSSA progenitor of ST772 on the Indian subcontinent.

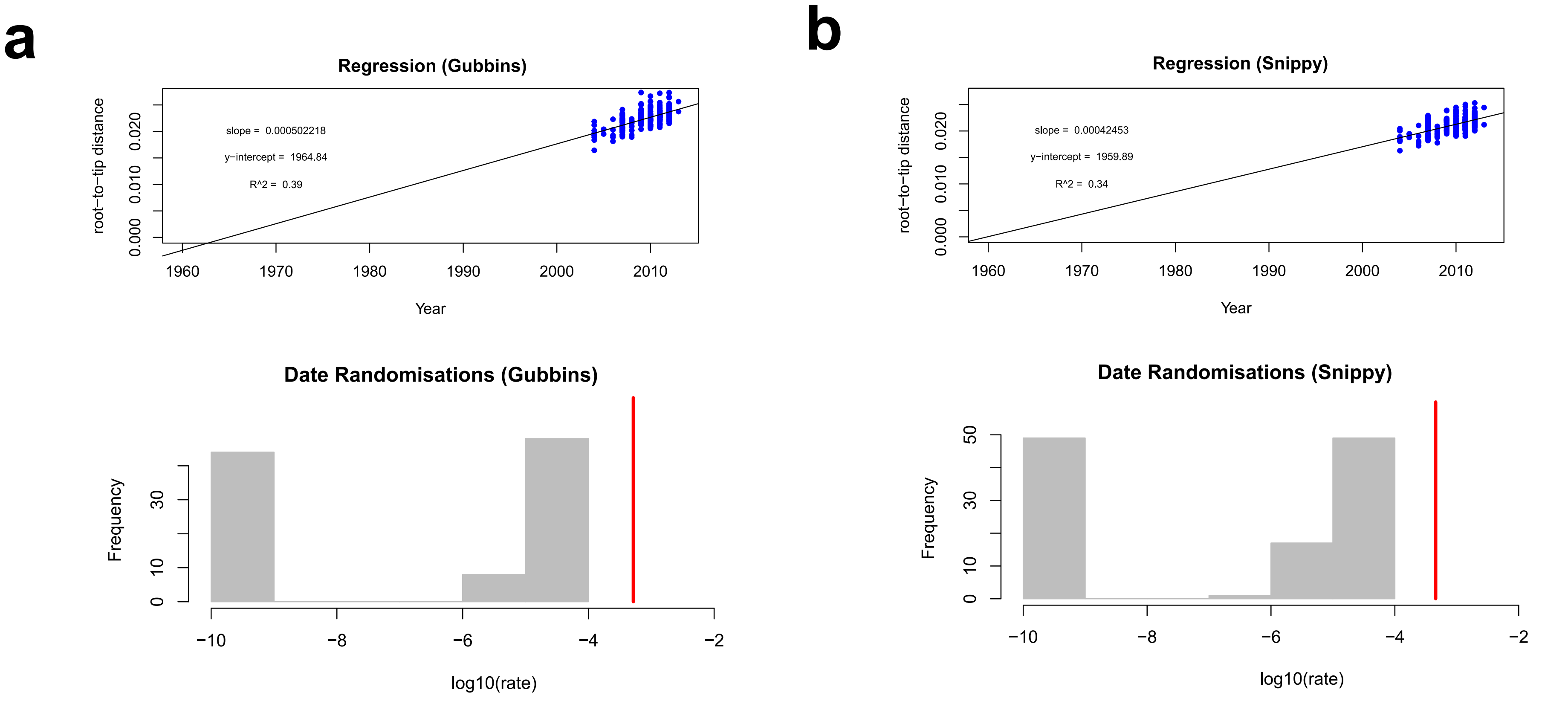


**Fig. S4:** Regression analysis and date randomisations for Least Squares Dating (LSD) analysis of ML phylogenies computed from core variants with PhyML under the GTR + G model (a) before (Snippy, 7,039 SNPs) and (b) after removing recombination (Gubbins, 7,928 SNPs). Each blue point in the regression plots corresponds to the distance from the root to a tip in the tree, and the solid black line is the least-squares regression. The substitution rate (slope), time to the most recent common ancestor (TMRCA, x-intercept), and $R^2$ (degree of clocklike behaviour) are shown. In the date randomisations, the red line denotes the substitution rate estimate from LSD using the correct sampling times. The gray bars correspond to the histogram of substitution rate estimates obtained by randomising the sampling times 100 times, such that they represent the null distribution of rate estimates under no temporal structure. The rate estimate with the correct sampling times is not within the range of those obtained using randomisations, indicating that the data have strong temporal structure.

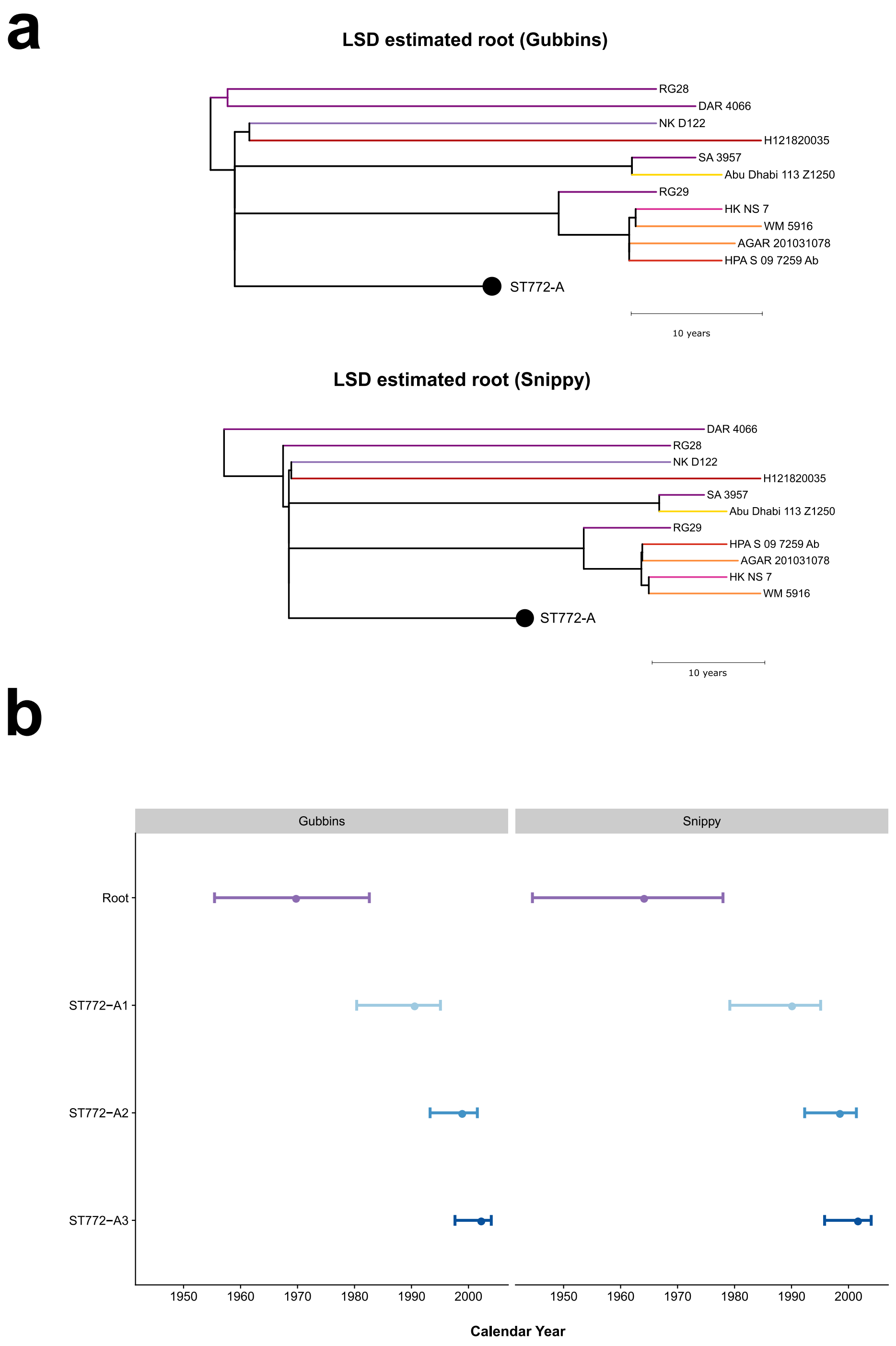


**Fig. S5:** (a) Rooting of maximum likelihood phylogenies computed from core variant alignments before (Snippy, 7,039 SNPs) and after removing recombination (Gubbins, 7,928 SNPs) optimised in Least Squared Dating (LSD) (see Fig. S4). (b) Divergence date estimates of the root and major population sub-groups in ST772 computed in LSD for ML phylogenies before (Snippy) and after removing recombination (Gubbins), including 95% confidence intervals for nodes (CI) using parametric bootstrapping in LSD.

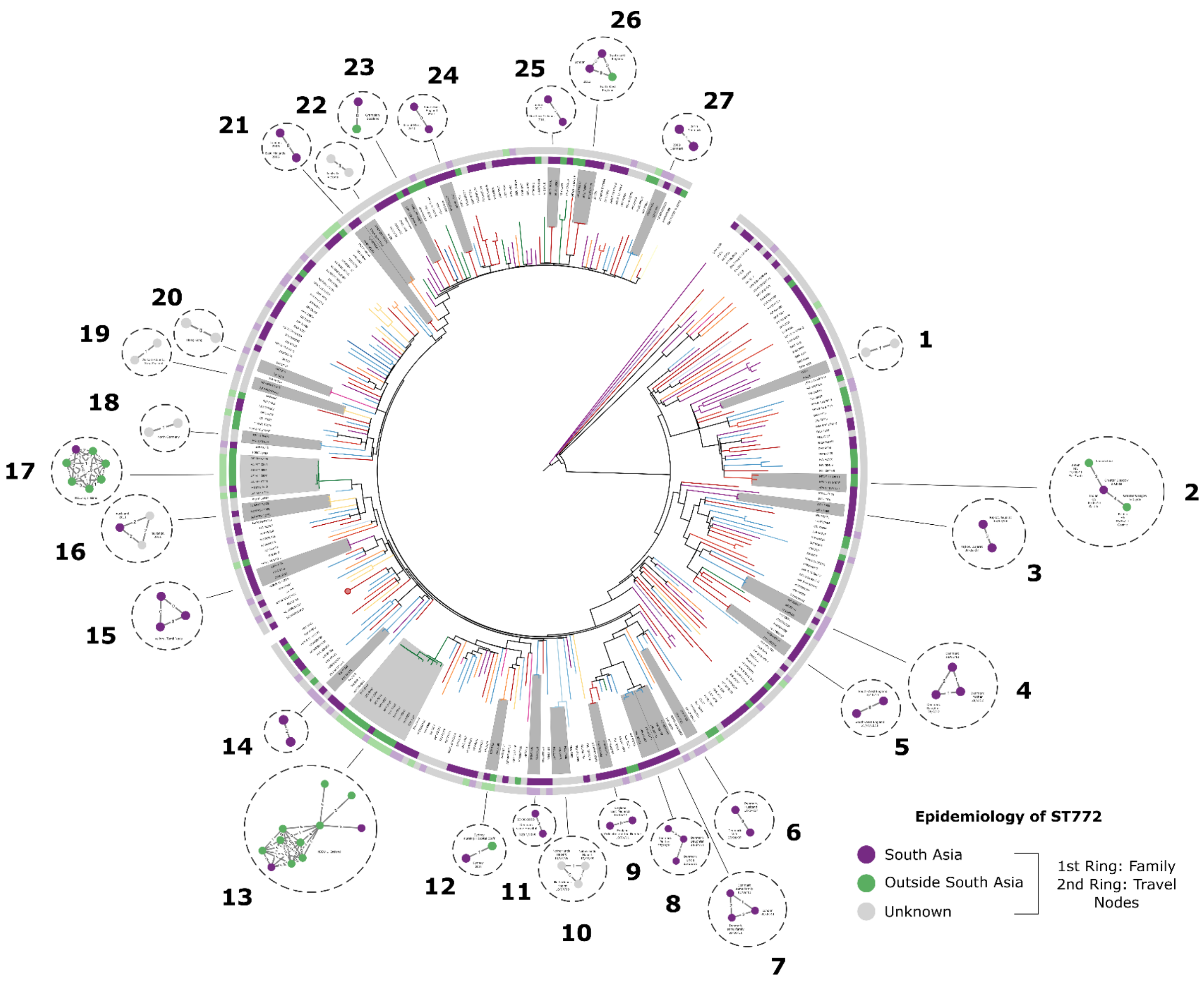


**Fig. S6**: Transmission clusters in ST772 (see also SI Table 10). Connected components of undirected graph with a threshold on pairwise SNP distance (4 SNPs, corresponding to maximum SNP distance within one year at core genome substitution rate of 1.61 x $10^{-6}$ nucleotide substitutions/site/year inferred with Least Squares Dating) and mapped to maximum likelihood phylogeny of ST772. Inner and outer ring depict family and travel links to South Asia (India, Pakistan, Nepal), while node colours within the dotted circles of clusters depict combined epidemiological link to South Asia (see Methods). Transmission clusters detected with network approach are highlighted in gray within the phylogeny of ST772. Table indicates notable epidemiological links. Most clusters (19/27) indicate a family or travel link to South Asia. Some of these cluster may indicate transmission within house-holds, for instance cluster 6 (husband and wife who both travelled to India) or cluster 7 and cluster 8, which include putative transmission between family members who also have family links to South Asia. Other clusters (cluster 2, 12, 23, 26) have at least one member with links to South Asia and another without links to the Indian subcontinent, indicating spread within the community. NICU-1 and NICU-2 denote outbreaks in neonate intensive care units in Ireland. In NICU-1, the suspected index case was a staff member (M11-0092) who had been hospitalized in India, where she had travelled shortly before the recovery of the first isolate and where she had given birth to her child (M11-0167). The child was screened after detection in the index staff member, following the first three neonate cases in 2010 (isolates M10). A high resolution image of the specific connected components and phylogeny in available in Fig. S2.

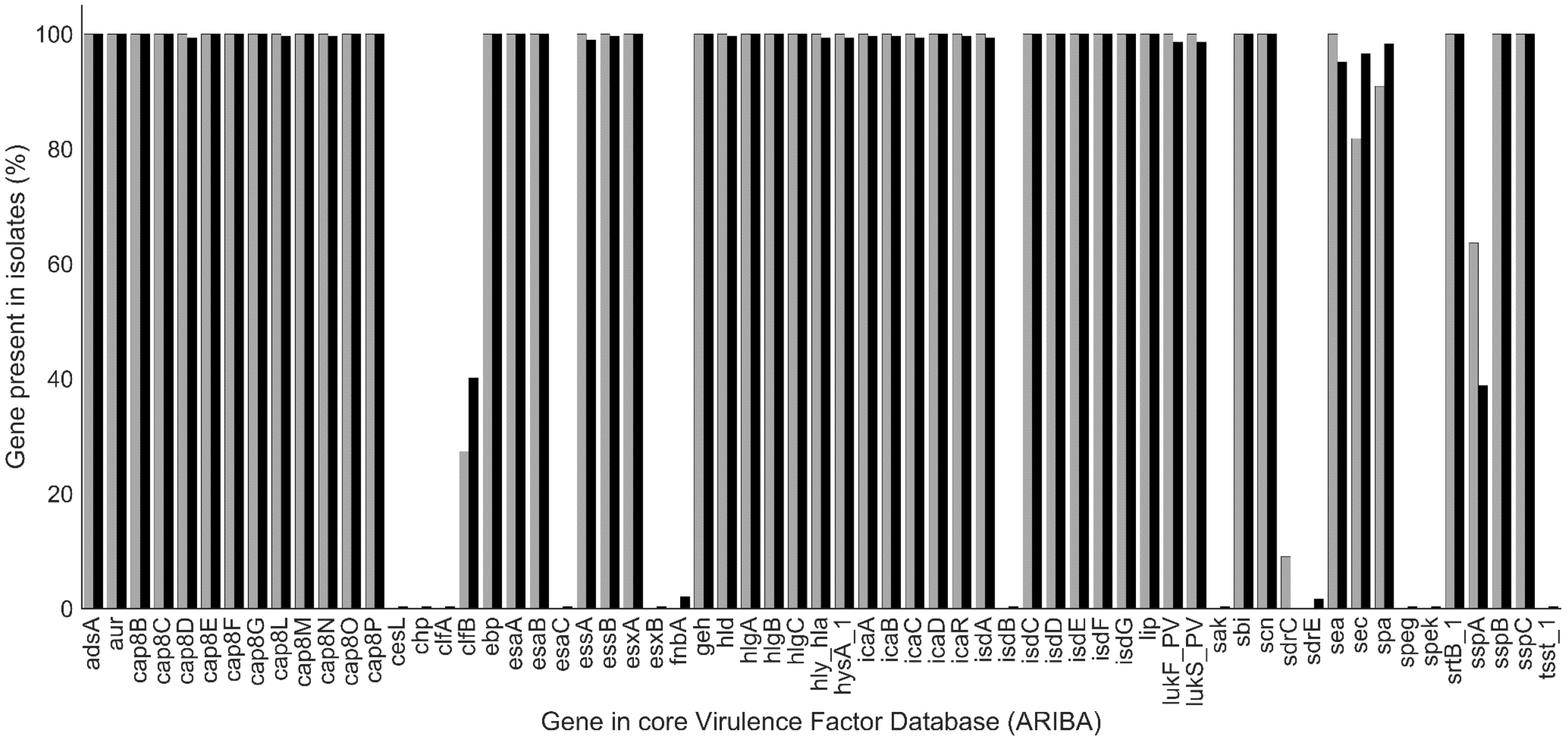


**Fig. S7:** Proportion of isolates in basal strains (gray) and ST772-A (black) carrying virulence factors from the core virulence factor database, detected with ARIBA. There are no significant differences in virulence factor carriage between basal strains and ST772-A at $p < 0.01$ using Fisher's exact test.

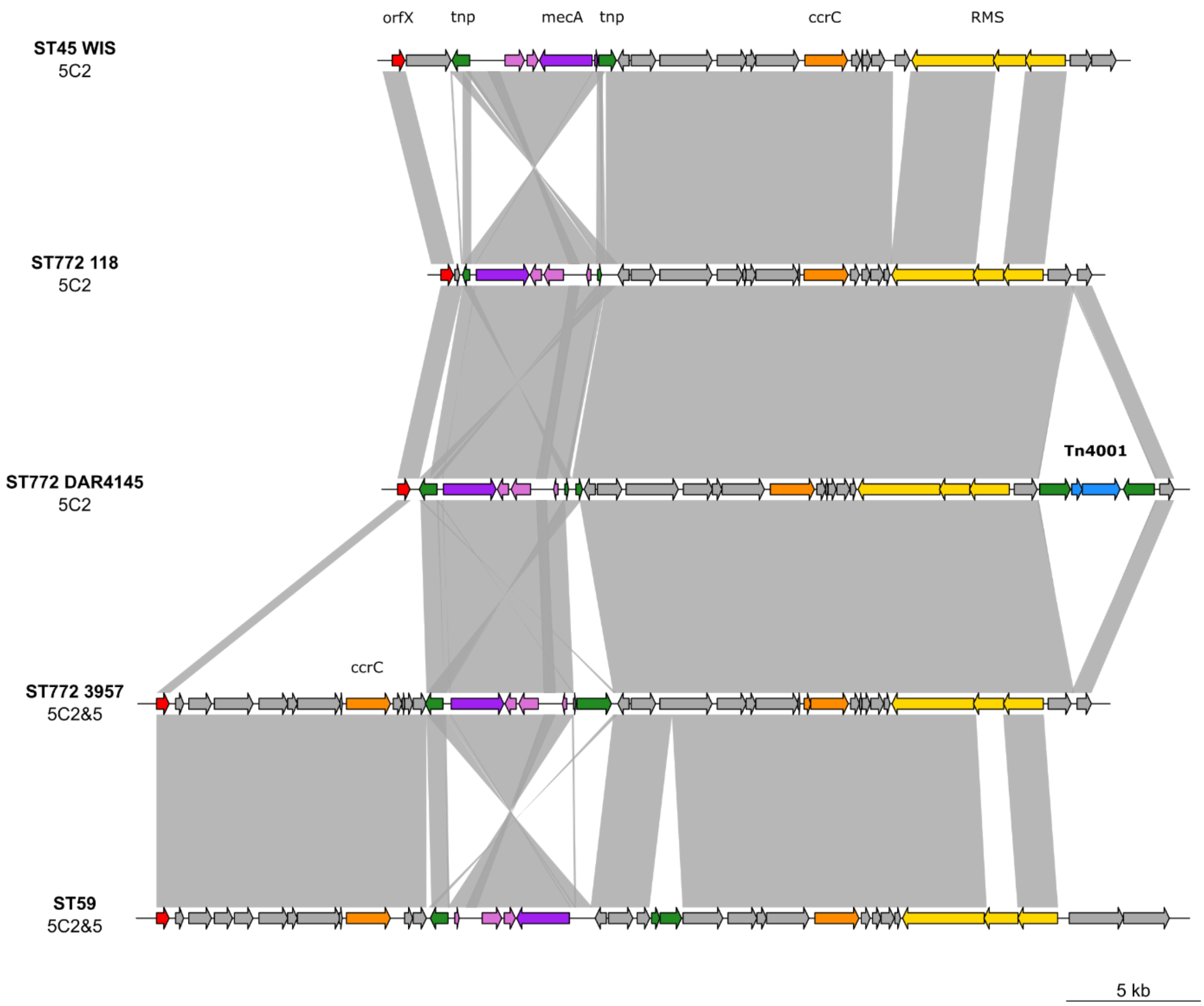


**Fig. S8:** SCC*mec* structure of main subtypes 5C2 and 5C2&5 present in ST772, including integration of the Tn*4001* transposon in subtype 5C2. Nucleotide BLAST comparison (polygons, minimum identity > 85%) of different subtypes of SCC*mec*-V. ST772 carries two subtypes, 5C2 and the composite cassette 5C2&5 downstream of *orfX* (red), which harbours an additional *ccrC* (orange). Both subtypes show the characteristic inversion of the *mec*-complex (purple, orchid), flanked by complete or truncated transposition elements (green). An additional major variant of 5C2 found in the reference genome DAR4145 harbours Tn*4001* downstream of the Type I restriction-modification system (yellow, RMS). The specificity sub-unit (middle, *hsdS*) of the RM system show < 85% nucleotide identity with other systems on SCC*mec*-V. This corresponds to unusual recognition site methylation in the ST772 PacBio reference genome DAR4145 (available at REBASE). Additional experiments will need to be conducted to match the recognition sites of all RM systems occurring in DAR4145 to the target recognition domains of *hsdS*.

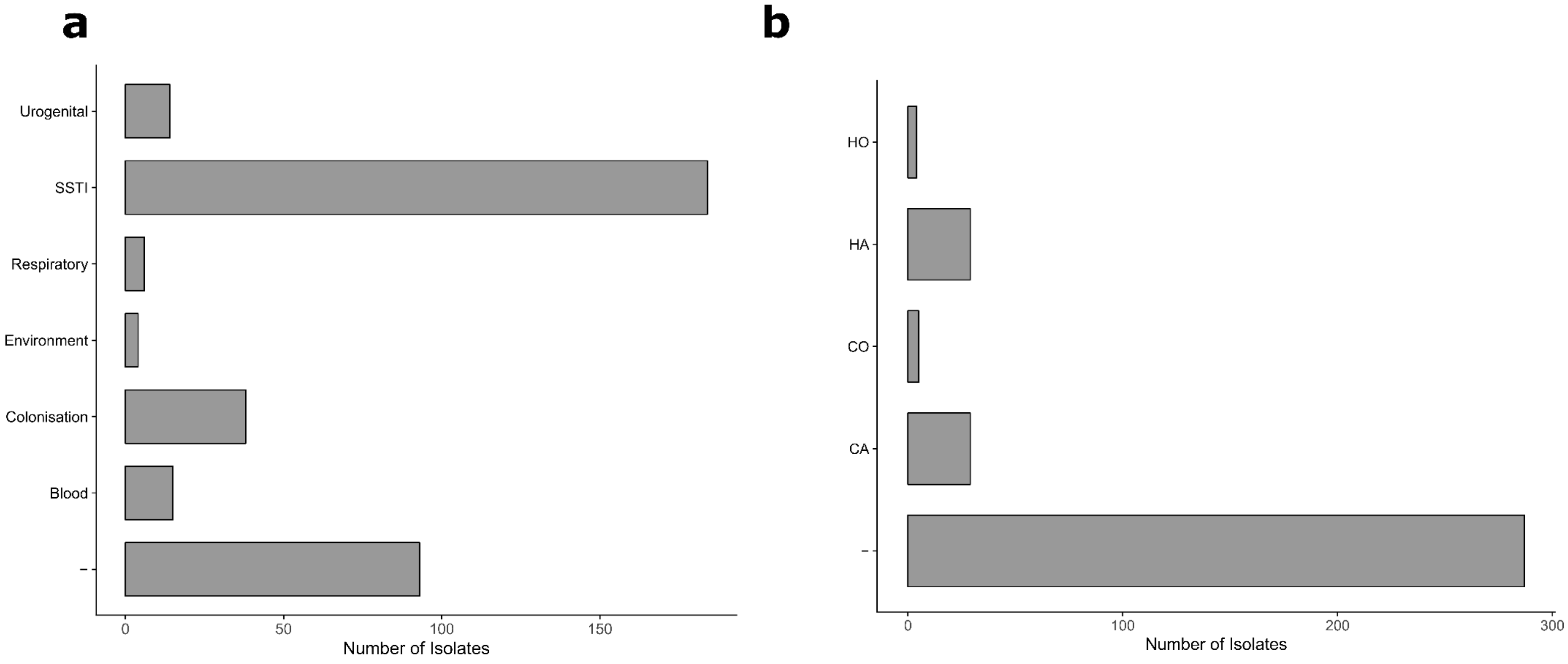


**Fig. S9:** Isolation and acquisition in ST772. (a) Source of isolation as described in Methods, data is available in Table S1. SSTIs and colonisation are the most prevalent presentation of ST772 (- = no data) (b) Acquisition of ST772 isolates isolation (HA = healthcare-associated, CA = community-associated, CO = community-onset, HO = healthcare-onset, - = no data). Data of acquisition status is sparse and further studies may endeavour to collect reliable epidemiological data to map the circulation of the Bengal Bay clone in healthcare and community environments.

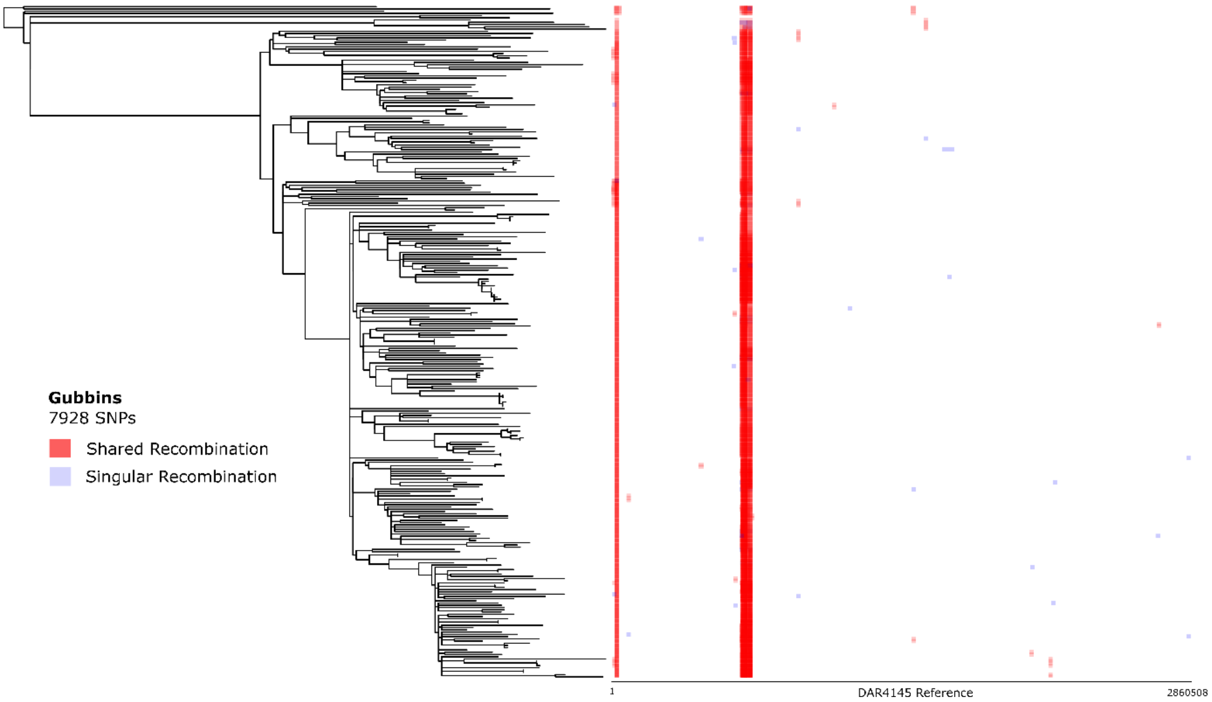


**Fig. S10:** Homologous recombination events detected by Gubbins across the ST772 phylogeny reconstructed from non-recombinant SNPs (205 recombinant segments, 7,928 SNPs). Longitudinal samples from a veterinary healthcare (VET, n= 39) are reduced to a single representative isolate. Red blocks indicate internal recombination (shared by common ancestry), while blue blocks indicate terminal recombination (singular occurrences). The majority of recombinant segments (74.6%) were detected in the core of a mosaic prophage as described in the reference genome DAR4145. Several segments were located within the SCC*mec* (8.8%), including one large (17,254 bp) segments spanning the *mec* complex and *ccrC*, shared between the cluster of 5 basal strains carrying SCC*mec-V* (5C2&5). This cluster also shared a 924 bp recombinant segment within the large extracellular matrix binding protein (*ebp*). Another notable occurrence was a small recombinant segment (501 bp) overlapping the 3' end of *ccrC* and an adjacent hypothetical protein in all isolates that harboured SCC*mec*. Other segments outside these regions included 40,019 bp of the PVL-prophage Phi-IND772 including *sea* and *lukF/S-PVL* in a single isolate from Scotland and two segments (3,387 and 4,197 bp) spanning several genes within the νSaα specific lipoprotein like cluster in one isolate from Hong Kong and two isolates from Denmark.

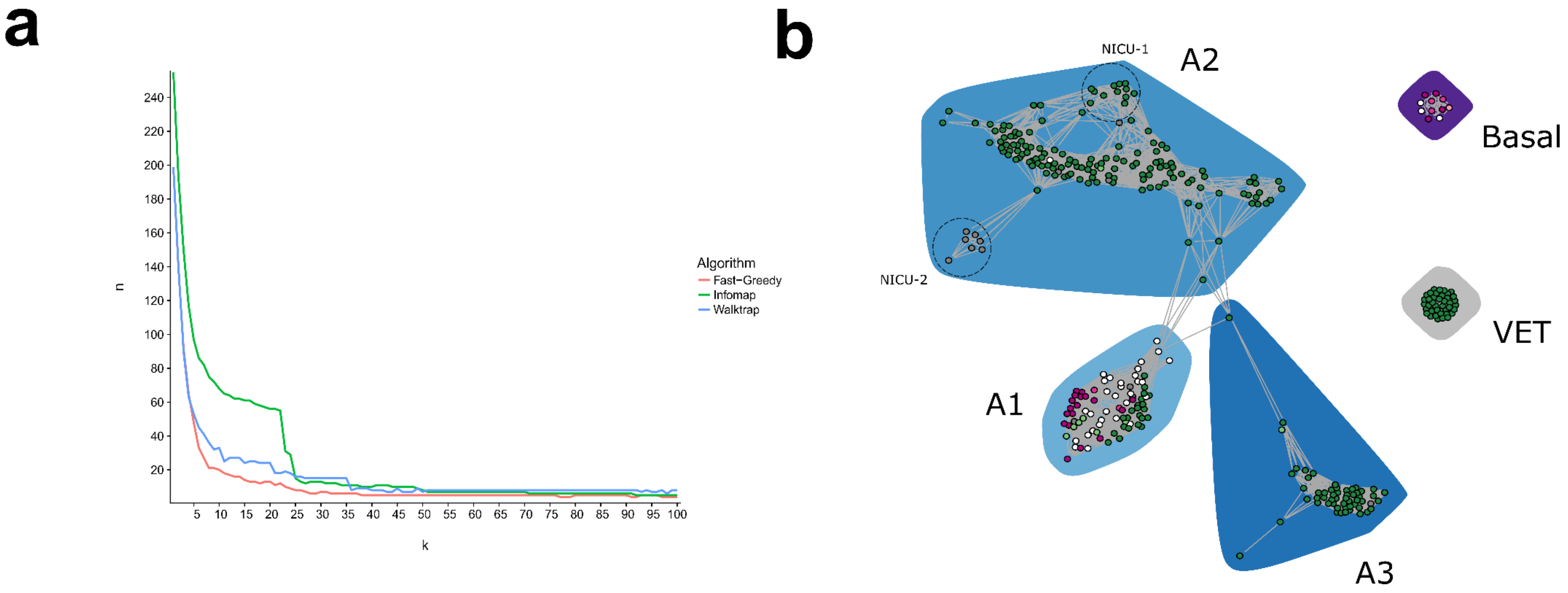


**Fig. S11:** Population network graph of ST772. (a) Plot of the number of detected communities (n) versus the number of mutual nearest neighbours (k) parameter in NetView (k = 1 - 100) using a pairwise Hamming distances and community detection algorithms as implemented in *igraph* and *NetView* v.1.1 (http://github.com/esteinig/netview). While community detection differs strongly in the assembly phase of the network (k = 1 – 25), the plot indicates a relative congruence of community detection between algorithms at k > 25-40. (b) Complete network topology (Fruchtermann-Reingold) at k = 40, showing communities from fast-greedy modularity optimisation (polygons) and SCC*mec* type as in Figure 1a. The veterinary cluster, excluded from the main network in Figure 1c, is shown in this representation (gray polygon, VET).

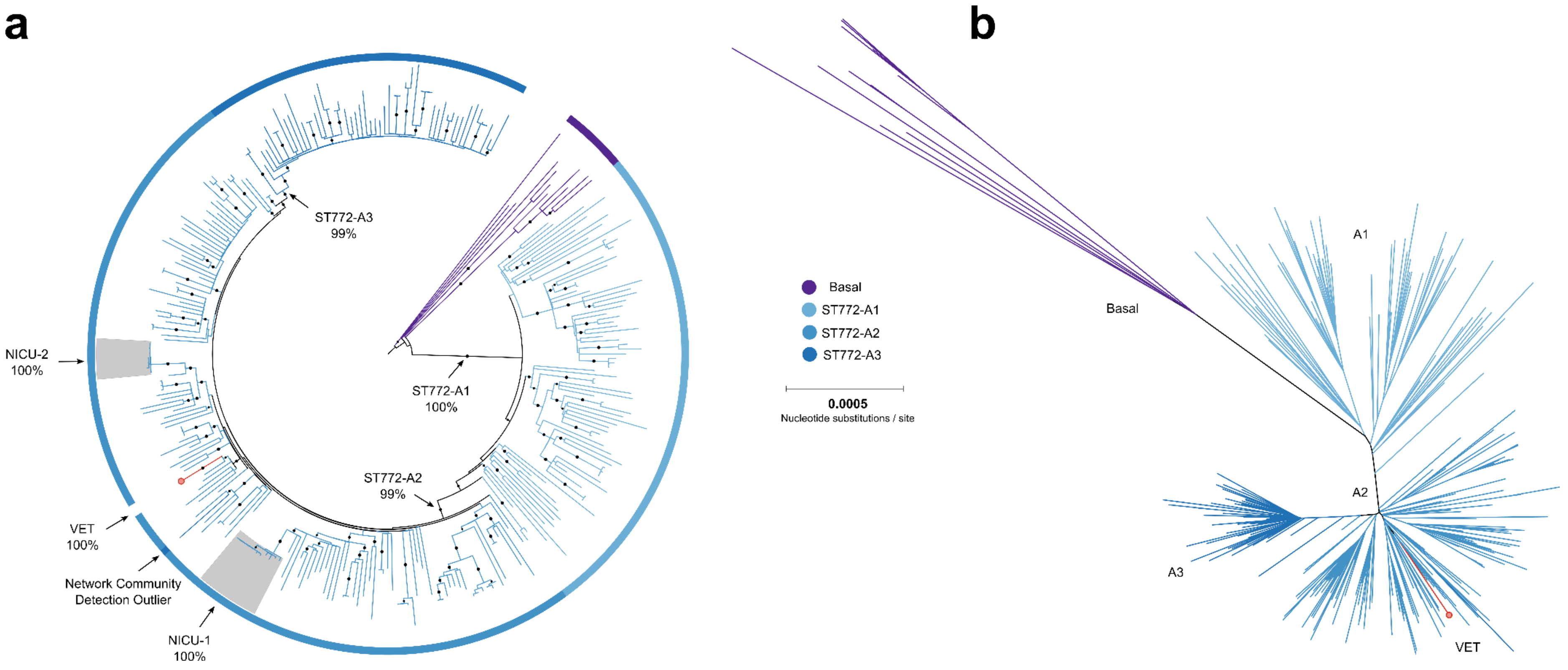


**Fig. S12**: Maximum likelihood phylogenies of ST772 showing community clusters detected with NetView (colors) and their bootstrap support values in (a) the mid-point rooted main phylogeny, with black circles indicating bootstrap support > 95% and as (b) unrooted phylogeny, demonstrating a pattern of rapid population divergence and proliferation of ST772-A

**a**

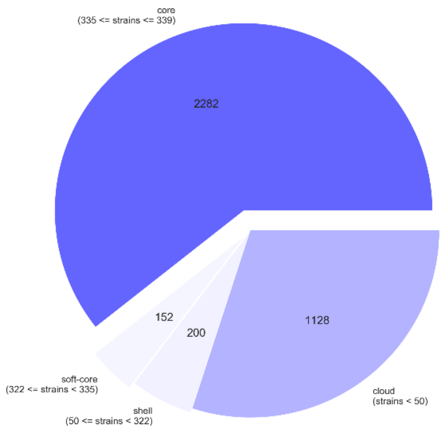


**b**

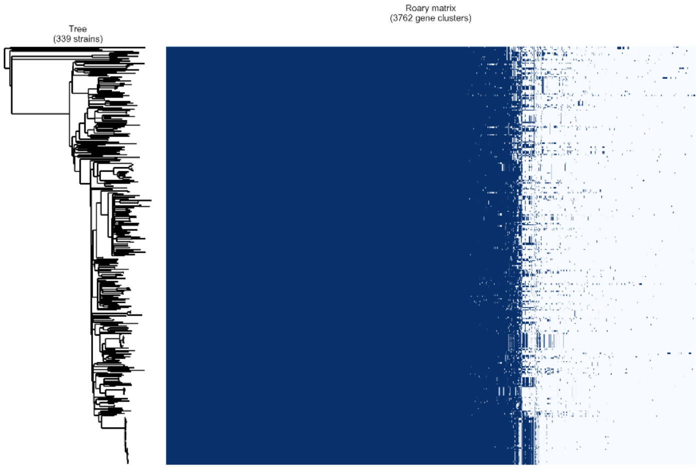


**Fig. S13:** Pan-genome of ST772 computed from assembled and Prokka-annotated genomes (n = 339, excluding reference genome DAR4145) using Roary. The lineage demonstrates a stable core genome (genes occurring in > 98% of strains) with little variation in the accessory genome. Pie chart and contiguous visualization were determined across the core genome phylogeny, using *roary_plots.py* (https://github.com/sanger-pathogens/Roary/contrib/roary_plots). Accessory genes comprise mainly phage-associated coding regions, plasmid-associated genes and specific genes rarely observed across the population of ST772, such as a type I restriction-modification system (RMS), lipoprotein like genes, cassette recombinases and fibronectin binding protein A (data available at the repository).
